# Structures of the Human LONP1 Protease Reveal Regulatory Steps Involved in Protease Activation

**DOI:** 10.1101/2020.11.23.394858

**Authors:** Mia Shin, Edmond R. Watson, Scott J. Novick, Patrick Griffin, R. Luke Wiseman, Gabriel C. Lander

## Abstract

The human mitochondrial AAA+ protein LONP1 is a critical quality control protease involved in regulating diverse aspects of mitochondrial biology including proteostasis, electron transport chain activity, and mitochondrial transcription. As such, genetic or aging-associated imbalances in LONP1 activity are implicated in the pathologic mitochondrial dysfunction associated with numerous human diseases. Despite this importance, the molecular basis for LONP1-dependent proteolytic activity remains poorly defined. Here, we solved cryo-electron microscopy structures of human LONP1 to reveal the molecular mechanism of substrate proteolysis. We show that, like bacterial Lon, human LONP1 adopts both an open and closed spiral staircase orientation dictated by the presence of substrate and nucleotide. However, unlike bacterial Lon, human LONP1 contains a second spiral staircase within its ATPase domain that engages substrate to increase interactions with the translocating peptide as it transits into the proteolytic chamber for proteolysis. Further, we show that substrate-bound LONP1 includes a second level of regulation at the proteolytic active site, wherein autoinhibition of the active site is only relieved by the presence of a peptide substrate. Ultimately, our results define a structural basis for human LONP1 proteolytic activation and activity, establishing a molecular framework to understand the critical importance of this protease for mitochondrial regulation in health and disease.

## Introduction

Mitochondria are the site of essential cellular functions including oxidative phosphorylation, apoptotic signaling, calcium regulation, and iron-sulfur cluster biogenesis. The maintenance of mitochondrial protein homeostasis (or proteostasis) is critical for mammalian cell viability^1,2^. To prevent the accumulation of misfolded, aggregated, or damaged proteins, mitochondria evolved a network of quality control proteases that function to regulate mitochondrial proteostasis in the presence and absence of cellular stimuli^1,3^. This includes the ATP-dependent protease LONP1 – a protease that is conserved throughout evolution from bacteria to humans^1,3–5^. In mammals, LONP1 is a primary quality control protease responsible for regulating proteostasis and function within the mitochondrial matrix through diverse mechanisms including the degradation of misfolded and oxidatively damaged proteins, components of the electron transport chain, and mtDNA regulatory factors such as TFAM and POLG^4–10^. The importance of LON in regulating mitochondria is underscored by the fact that homozygous deletion of *Lonp1* in mice is embryonic lethal^11^. Further, imbalances in LONP1 activity are implicated in mitochondrial dysfunction associated with diverse pathologic conditions including organismal aging, cancer, and numerous other human diseases^1,4,11–14^. Mutations in *LONP1* have also been identified to be causatively associated with cerebral, ocular, dental, auricular, skeletal (CODAS) syndrome, a multi-system developmental disorder consisting of a wide array of clinical manifestations including hypotonia, ptosis, motor delay, hearing loss, postnatal cataracts, and skeletal abnormalities^15–17^. Despite the central importance of LONP1 for regulating mitochondria in health and disease, the structural basis for human LONP1 proteolytic activation and activity remains poorly defined.

The LONP1 protease consists of an N-terminal domain involved in substrate recognition and oligomerization, a AAA+ (**A**TPases **a**ssociated with diverse cellular **a**ctivities) domain that that powers substrate translocation, and a C-terminal serine protease domain involved in substrate proteolysis^1,4,18^. Numerous biophysical approaches have been employed to gain insight into the macromolecular structure and function of bacterial Lon^19–27^, and recent cryo-electron microscopy (cryo-EM) studies of bacterial Lon in both substrate-free and substrate-bound conformations revealed the structural basis for substrate processing and protease activation^28^. However, few structural studies have been directed towards understanding the molecular mechanism of protease activation and substrate processing by human LONP1. A prior crystal structure of the isolated protease domain of human LONP1 (PDB:2X36) revealed the structural conservation between human and bacterial Lon protease domains, suggesting similar mechanisms of action^29^. Further, a low resolution cryo-EM structure of human LONP1 showed the ATPase and protease domains organized to form a pseudo six-fold symmetric chamber, while the N-terminal domains are arranged as a trimer of dimers atop the AAA+ domains^30^. Given the complex proteolytic demands of the mitochondrial matrix, we aimed to define the structural and mechanistic differences from bacterial Lon that have evolved to improve the activity or regulation of this protease for mitochondrial regulation.

To address this deficiency, we solved cryo-EM structures of human LONP1 in the presence and absence of a translocating substrate. We show that in the absence of substrate, LONP1 adopts an open, left-handed spiral conformation that is bound to ADP and has its protease active sites organized in an auto-inhibited conformation. In contrast, in the presence of substrate, the AAA+ domains of human LONP1 adopt a right-handed spiral configuration similar to that observed for many other AAA+ proteases^18,31,32^. Surprisingly, and unlike other structures of AAA+ proteases, including bacterial Lon^28^, we found that the protease domains in substrate-bound human LONP1 adopt an asymmetric arrangement wherein the protease active sites remain auto-inhibited. This asymmetric arrangement of the protease domains could be relieved by administration of the small molecule LONP1 inhibitor bortezomib, which covalently engages the proteolytic active site and gives rise to the six-fold symmetric arrangement of active protease domains, similar to that observed for bacterial Lon^28^. This bortezomib-induced release of LONP1 protease auto-inhibition suggests a second level of regulation for protease activation that has evolved to tune LONP1-dependent regulation of mitochondrial proteostasis and function.

## Results

### Substrate-bound and substrate-free structures of human LONP1

Mature human LONP1 lacking its N-terminal mitochondrial targeting sequence was incubated with saturating amounts of ATPγS (1 mM), a slowly hydrolyzing ATP analog, to stabilize the complex for structural studies. Single-particle cryo-EM analyses resulted in reconstructions of two distinct conformations of the essential protease: one structure containing a peptide substrate trapped in its central channel and the other devoid of substrate (**Fig. 1** and **Supplementary Fig. 1**). The presence of substrate in a subset of the complexes was attributed to co-purification of an endogenous protein substrate, as has been observed previously in cryo-EM studies of numerous other AAA+ proteins^18,31–37^. In both structures of human LONP1, residues 420-785, comprising a portion of the N-terminal helical domain (NTD^3H^), the AAA+ domain consisting of the small and large ATPase subdomains, and the protease domain, were well-resolved (**Fig. 1**); however the entirety of the 300-residue N-terminal domain was not resolvable in either structure, likely due to the inherent flexibility of this region.

**Figure 1.**
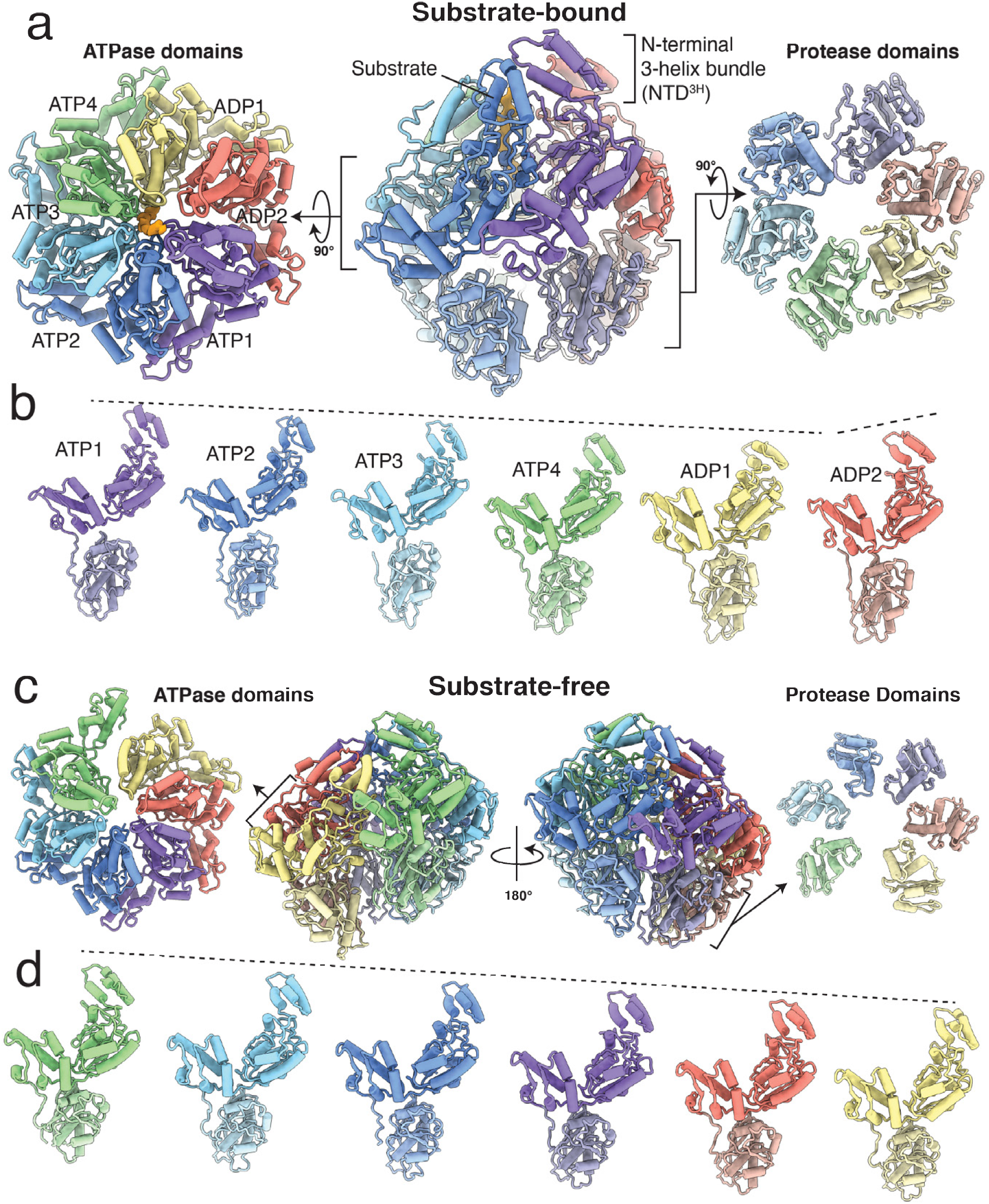
Architectures of substrate-bound and substrate-free human mitochondrial LONP1 protease. **a.** Lateral view of the substrate-bound human LONP1 atomic model in a left-handed spiral configuration (center) flanked by axial exterior views of the ATPase (left) and protease (right) domain rings. Each subunit of the homohexamer is assigned a distinct color depending on its position in the spiral staircase, with the protease domains colored a lighter hue than the ATPase domains. The same coloring is used throughout the figure. The cryo-EM density of the substrate is shown as a solid isosurface colored orange. **b.** The relative height of individual protomers within the substrate-bound conformer are shown by aligning all subunits to a common view, and accentuated by dashed lines shown above the NTD^3H^. **c.** The same orthogonal views in (**a**) are shown for the substrate-free human LONP1 atomic model. Each subunit of the homohexamer is assigned a distinct color depending on its position in the spiral staircase, **d.** The individual protomers of substrate-free human LONP1 lined up side-by-side, showing the descending positions of the domains in the structure.

The substrate-free human LONP1 reconstruction identified in our dataset was resolved to 3.4 Å resolution, which was sufficient for atomic modeling for five of the six LONP1 protomers (**Supplementary Fig. 1** and **Supplementary Table 1**). The organization is reminiscent of the left-handed, open lockwasher configurations previously observed for substrate-free bacterial Lon^26,28^, although human LONP1 appears to oligomerize with a slightly smaller helical pitch than the bacterial forms (**Fig. 1c,d** and **Supplementary Fig. 2**). The uppermost subunit of substrate-free human LONP hexamer demonstrates substantially more flexibility than its bacterial counterpart, as it is only discernible at low resolution in a subset of the particles (**Supplementary Fig. 1**), suggesting that oligomerization of the N-terminal domains into a trimer-of-dimers^30^ may play a role in maintaining the hexamer’s oligomeric state in the absence of substrate. Due to the low resolution of this uppermost subunit, a copy of the middle protomer from the five-subunit spiral was rigid body fit into the reconstruction for representational purposes. Similar to previous substrate-free structures of bacterial Lon^26,28^, the subunits of human LONP1 in the absence of substrate are bound to ADP and all protease subdomains adopt an auto-inhibited conformation (**Supplementary Fig. 3**). This substrate-free structure likely represents a conformation important for substrate engagement and release, as suggested for bacterial Lon^28^. According to this mechanism, the topmost subunit of the left-handed spiral is the first to undergo nucleotide exchange and engage substrate, followed by subsequent nucleotide and substrate engagement events that lead to a reorganization of the AAA+ domains within the hexamer into an active state that is competent for substrate translocation. The similarity of the substrate-free human and bacterial structures indicates that this model for substrate engagement is conserved.

The substrate-bound LONP1 structure was resolved to 3.2 Å resolution, which enabled atomic model building and refinement (**Supplementary Fig. 1** and **Supplementary Table 1**). The structural conservation of the ATPase domains from bacteria to human is considerable, as the protomers of the human LONP1 are in a nearly identical organization as their counterpart in *Y. pestis* Lon, with RMSDs between 0.8-0.9 Å (**Supplementary Fig. 4a**). The overall organization of these ATPase domains in the substrate-bound human LONP1 generally resembles that of previously determined substrate-bound AAA+ proteins, wherein the ATPases form a closed spiral staircase around a centrally positioned substrate peptide.

However, our substrate-bound human LONP1 exhibits several notable structural differences from the previously determined substrate-bound *Y. pestis* Lon. Whereas the AAA+ ring of bacterial Lon consists of four descending ATPase domains in its spiral staircase with two ascending ‘seam’ subunits (**Supplementary Fig. 5**), our substrate-bound structure of human LONP1 contains five descending ATPase domain subunits with a single ‘seam’ subunit transitioning between the highest and lowest subunits (**Fig. 1a,b**). This conformation is consistent with substrate-bound structures of AAA+ protein translocases involved in a wide range of biological functions from bacteria to mammals^18,32–37^. Accordingly, the four topmost subunits of the staircase are bound to ATPγS (hereafter referred to as ATP for simplicity) while the lowest contains ADP in its nucleotide binding site (**Fig. 1a,b** and **Supplementary Fig. 6**). Density corresponding to an ADP molecule is also observed in the ‘seam’ subunit. To maintain consistency with other AAA+ proteins, we will name the ATP-bound subunits, starting with the topmost subunit that is bound to ATP, ATP1-4, the lowest subunit in the spiral staircase that is ADP-bound will be referred to as ADP1, and the ‘seam’ subunit as ADP2 (**Fig. 1b** and **Supplementary Fig. 6**).

### Multiple pore loop interactions facilitate substrate translocation in human LONP1

The distinct configuration of the ATPase domains observed in human LONP1, relative to bacterial Lon, notably influences interactions with the engaged substrate. AAA+ proteins engage translocating substrates through pore loop residues within the ATPase domain that extend toward the central channel of the ATPase ring^18^. Previous structures of substrate-engaged AAA+ proteins have shown that aromatic residues in the ATPase pore loops intercalate into the backbone of an engaged peptide substrate every two amino acids to facilitate substrate translocation^18^. The ATPases in our human LONP1 reconstruction encircle a density corresponding to a 12-residue peptide substrate, which, surprisingly, is nearly twice the length of engaged substrate visualized in *Y. pestis* Lon^28^. This observation provides an opportunity to examine the evolutionary differences in substrate translocation by Lon proteases in bacteria and human.

In human LONP1, the substrate-interacting Y565 in pore loop 1 from all four ATP-bound subunits, ATP1-4, as well as the ADP-bound subunit ADP1 at the bottom of the spiral staircase, are all observed engaging with the translocating substrate (**Fig. 2a,b**). This is in contrast to the substrate-interacting arrangement previously shown for bacterial Lon, where only the three uppermost ATP-bound subunits were tightly engaged with substrate, while the fourth subunit within the spiral only weakly interacts with substrate^28^. Instead, the substrate interactions we observe in human LONP1 more closely resemble other substrate-bound structures of AAA+ proteins than those in bacterial Lon^18,31,32^.

**Figure 2.**
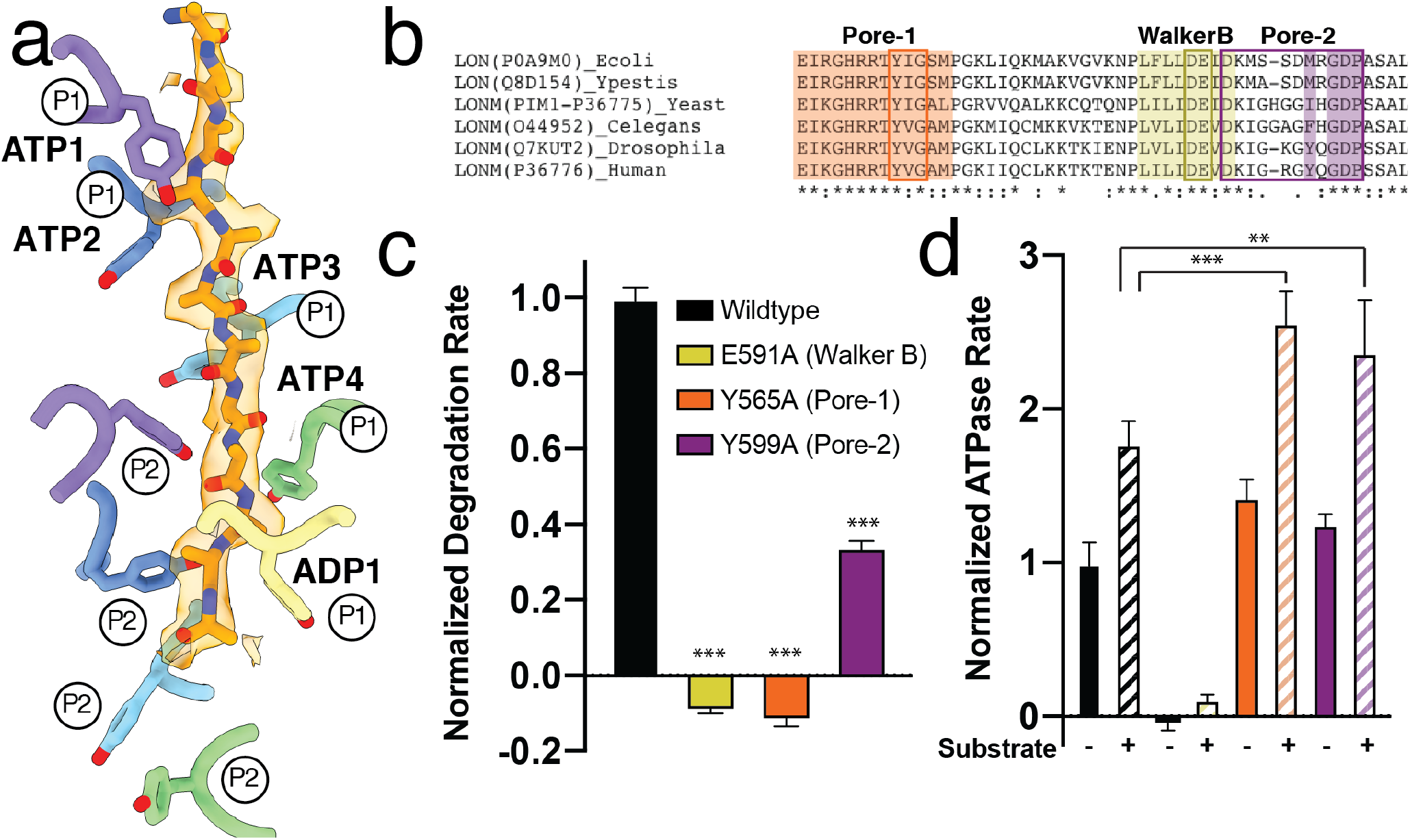
LONP1 employs two pore loop aromatic residues to facilitate substrate translocation. **a.** A twelve-residue polyalanine chain is modeled into the substrate density found in the substrate-bound human LONP1 structure, shown in a transparent orange surface representation. Y565 from pore-loop 1 (labeled P1) is shown using stick representations. Y565 residues from ATP1-4 and ADP1 subunits show intercalating, zipper-like interactions with substrate. Y599 from pore loop 2 (labeled P2) also shows intercalating interactions with substrate in the ATP1 (purple) and ATP2 (dark blue) subunits. **b.** Sequence alignment between LONP1 homologs highlighting evolutionary conservation within pore loop 1 and Walker B motifs, as well as the evolutionary integration of a hydrophobic residue into pore loop 2 in metazoans. **c.** Introducing pore loop mutations Y565A or Y599A shows a decrease in the degradation rate of a model substrate, FITC-casein. The hydrolysis-blocking Walker B mutation E591A is included as a control. Error bars show standard deviation from 3 independent replicates. ***indicates p<0.001 relative to wildtype calculated by ANOVA. **d.** Basal-level (filled bars) and casein-stimulated (hatched bar) ATPase rates are shown for the same mutations as in (**c**). Error bars show standard deviation from 3 independent replicates. **indicates p<0.01 and ***indicates p<0.001 relative to wildtype calculated by ANOVA.

Notably, in addition to the pore loop 1 aromatic residue, we identified a second pore loop aromatic residue, Y599, within the AAA+ domains of human LONP1 that also engages the translocating substrate (**Fig. 2a**). This second pore loop residue assembles into a spiral that parallels the pore loop 1 aromatic, with pore loop 2 from ATP1 and ATP2 engaged with the trapped substrate (**Fig. 2a**). While the remaining subunits do not appear to engage substrate in our structure of human LONP1, the spiraling organization of these residues and their proximity to translocating substrates suggest that these subunits may facilitate substrate guidance through the central channel and into the proteolytic chamber. Interestingly, the identified interactions between the pore loop 2 aromatics of ATP1 and ATP2 and the translocating substrate extend the overall interactions between human LONP1 and substrate deeper into the central chamber. The presence of a methionine, which cannot establish intercalating interactions with substrate, at the equivalent residue in *Y. pestis* Lon (M432), is likely responsible for a lack of observable pore loop 2 interactions in its structure^28^ (**Fig. 2b** and **Supplementary Fig. 7**).

Consistent with an important role for the pore loop 2 residue Y599 in substrate translocation in human LONP1, recombinant LONP1 containing a Y599A mutation exhibited changes in both substrate proteolysis and ATP hydrolysis. LONP1^Y599A^ showed a 66% decrease in the degradation rate of the model substrate FITC-casein (**Fig. 2c**) while showing a 36% increase in substrate-induced ATPase hydrolysis (**Fig. 2d**). This decreased degradation coupled with increased ATP consumption suggests that the Y599A mutant is a less efficient translocase compared to the wildtype enzyme, with increased likelihood of substrate “slippage” and unproductive ATP hydrolysis, as previously proposed for other AAA+ proteins^38^. Similar results were observed with the pore loop 1 mutant LONP1^Y565A^, indicating that these two pore loop residues are primarily important for substrate processing. Thus, inclusion of an aromatic residue, capable of intercalating into the backbone of an incoming substrate, into the pore loop 2 of human LONP may have evolved to increase interactions with the substrate to improve translocation and degradation efficiency, as well as to impart a degree of substrate specificity, within the mitochondria. Other AAA+ translocases have been shown to utilize multiple pore loops to facilitate translocation, where it has been speculated that the integration of additional aromatic-containing pore loops into the translocation mechanism increases the ATPase motor’s ‘grip’ on an incoming substrate^18,31,32,34,36,39^.

### Substrate-bound human LONP1 adopts an auto-inhibited protease conformation

Another striking structural divergence of our substrate-bound structure of human LONP1 from bacterial Lon is in the organization of the protease domains. Previous results from bacterial Lon showed that substrate binding in four ATPase subunits and one ATP hydrolysis event promotes a rearrangement of the protease domains into a six-fold symmetric configuration that releases the proteolytic active site from an auto-inhibited conformation, inducing the formation of a binding groove to position substrates for proteolysis^28^. Surprisingly, despite having a peptide substrate tightly bound within the channel, the protease domains of substrate-bound human LONP1 do not adopt a six-fold symmetric configuration (**Figs. 1a and 3a**). Instead, the protease domains of the substrate bound hexamers follow the shallow spiraling trajectory of the AAA+ domains, with an opening in the proteolytic ring between ATP1 and ATP2, the two uppermost subunits of the spiral staircase (**Figs. 1b and 3a**). Further, we observe all six protease active sites to be in an auto-inhibited configuration, wherein a loop containing the catalytic serine residue (S855) is folded into a 3_10_ helix, sterically blocking access of substrates to the catalytic active site, while simultaneously an aspartic acid (D842) prevents the formation of the serine (S855)-lysine (K898) catalytic dyad, hindering substrate degradation (**Fig. 3b-e**).

**Figure 3.**
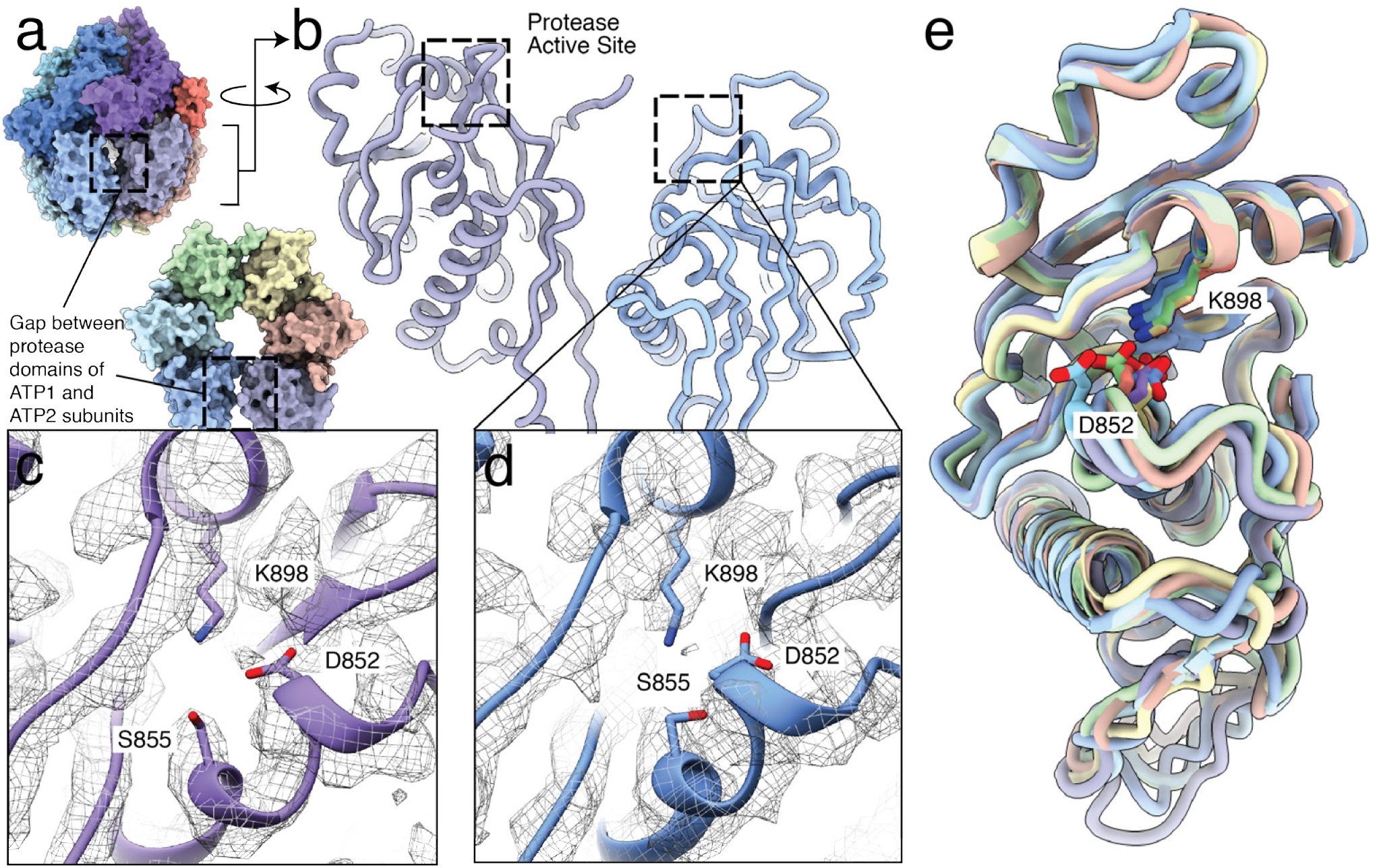
LONP1 in the substrate-bound conformation contains an autoinhibited protease site. **a.** Surface representation of substrate-bound LONP1 showing a break in the protease domains between the highest subunit, ATP1, and its neighboring subunit, ATP2, highlighted using a dashed box. **b.** The protease domains from ATP1 and ATP2 are shown using a ribbon representation, with the auto-inhibited active sites, denoted by dashed boxes. **c,d.** Details views of the regions denoted in (**b**) are shown using a ribbon representation within a mesh representation of the EM map. A loop containing the catalytic serine (S855) is folded into a 3_10_ helix and an aspartic acid (D842) prevents the formation of the serine (S855)-lysine (K898) catalytic dyad. **e.** An alignment of the isolated protease domains from all six subunits in substrate-bound LONP1 shows that all protease domains adopt an auto-inhibited conformation.

Previous results for bacterial Lon and other AAA+ proteases suggested that a flexible interdomain linker allows for large scale conformational rearrangements of the AAA+ domains independent of the symmetric protease to facilitate substrate translocation^28,33,40^. It was proposed that when the protease domains were positioned laterally alongside one another underneath the translocation-competent ATPase ring, the catalytic serine-containing loop would extend toward the neighboring protease domain to stabilize inter-subunit interactions, resulting in the formation of the six-fold symmetric protease ring with six active proteolytic sites^28,41^. However, even though several of the protease domains in the substrate-bound human LONP1 structure position alongside one another in an identical fashion as the those observed in the substrate-bound bacterial Lon structure, the catalytic loop remains in an auto-inhibited conformation (**Supplementary Fig. 8**). Thus, our structure of substrate-bound human LONP1 suggests that the combination of a translocating substrate and lateral positioning of protease domains is insufficient to produce the six-fold symmetric activated protease conformation observed in bacterial Lon, indicating other factors must be involved in generating a proteolytically active enzyme.

### Bortezomib relieves the protease auto-inhibition in substrate-bound human LONP1

One potential explanation for the lingering auto-inhibition observed in the protease active site of substrate-bound human LONP1 might be that, unlike bacterial Lon, substrate must be present to release the auto-inhibition of the protease active sites. To test this prediction, we determined a cryo-EM structure of substrate-bound human LONP1 incubated with saturating amounts of ATPγS as before, but further incubated with a ten-fold molar excess of bortezomib, a covalent, peptidomimetic inhibitor of LON that engages the proteolytic active site^6,25^. This complex, which we term LONP1^Bz^ to indicate the addition of bortezomib, was resolved to 3.7 Å resolution (**Supplementary Fig. 9** and **Supplementary Table 1**). Densities for the bortezomib molecules were clearly visible in each of the six active sites, positioned within the substrate binding groove and covalently attached to the catalytic serine (**Fig. 4a,d**).

**Figure 4.**
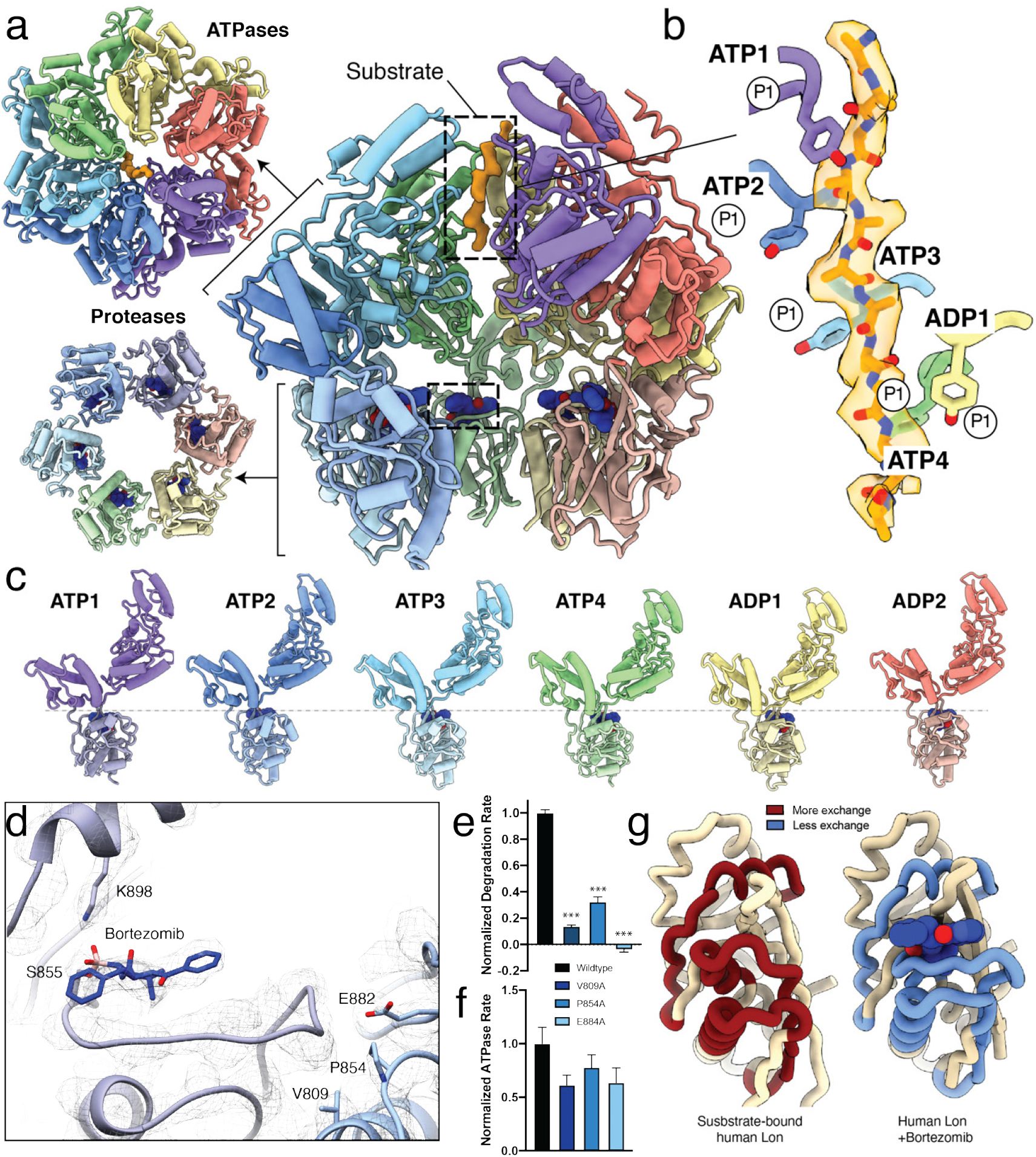
Bortezomib induces a six-fold symmetrization of the LONP1 protease domain and promotes activation of the proteolytic active site. **a.** Two external axial views of the bortezomib- and substrate-bound human LONP1Bz atomic model (left) is shown next to a cutaway lateral view of LONP1, using the same representation and coloring as in **Fig. 1**. Bortezomib (blue) bound within the protease active sites are shown using a sphere representation, with one bortezomib molecule denoted by a dashed box. **b.** A twelve-residue polyalanine chain is modeled into the substrate density within the central channel of the LONP1Bz structure, shown as in **Fig. 2**. The pore loop 1 Y565 residues from subunits ATP1-4 maintain intercalating, zipper-like interactions with substrate; however, Y565 from ADP1 in LONP1Bz is positioned further from substrate as compared to our substrate-bound structure (see **Fig. 2**). **c.** When the subunits are positioned side-by-side, the protease domains are aligned along a single plane. **d.** Bortezomib covalently attached to S855 in ATP1 is shown using a ribbon representation within a mesh representation of the cryo-EM map, where the loop containing the catalytic serine S855 is observed extended toward the neighboring protease domain and interacting with conserved residues V809, P854, and E882 from the neighboring protease domain, show in blue. **e, f.** Graphs showing rates of FITC-casein degradation (**e**) or ATPase rate (**f**) for LONP1 harboring alanine mutations in the conserved residues V809, P854, and E882 involved in stabilizing the catalytic loop in the active conformation of the protease active site. Error bars show standard deviation from 3 independent replicates. ***indicates p¡0.001 relative to wildtype calculated by ANOVA. **g.** A single protease domain is shown as a worm representation from the substrate-bound LONP1 structure in the absence (left) and presence (right) of bortezomib. Regions showing a decrease in D_2_O exchange measured by HDX-MS upon bortezomib binding are colored red in the auto-inhibited state (signifying greater exchange) and blue in the bortezomib-bound state (signifying decreased exchange).

Strikingly, bortezomib association with the protease active sites led to a dramatic reorganization of the LONP1 sub-units, resulting in a conformation that strongly resembles the substrate-bound bacterial Lon. The LONP1^Bz^ conformation contains four ATPγS-bound AAA+ domains arranged in a spiral, with two ADP-bound AAA+ domains adopting displaced ‘seam’ orientations (**Fig. 4a-c** and **Supplementary Fig. 10**). Despite differences in the spiral orientation of substrate-bound human LON and LONP1^Bz^, both structures show a similar engagement of a 12-residue translocating peptide by pore loop 1 aromatics (**Fig. 4b**). Intriguingly, we were unable to determine the interactions between pore loop 2 aromatics and substrate in our LONP1^Bz^ structure, as these loops were not as well-ordered as they were in the substrate-bound LONP1 structure, suggesting a dissociation of substrate from these pore loop 2 residues occurs prior or after substrate association with the protease domain.

Importantly, the protease domains in LONP1^Bz^ are assembled into a six-fold symmetric organization. Further, the presence of bortezomib relieved the auto-inhibition of the protease active site in our substrate bound LONP1, allowing proper assembly of the proteolytic active site, as previously observed for the substrate-bound bacterial Lon (**Fig. 4d** and **Supplementary Fig. 11**). This suggests that the presence of a substrate in the protease active site induces rearrangement of the serine-containing catalytic loop, such that it extends toward the neighboring protease domain to allow formation of the proteolytically active S855-K898 catalytic dyad. Previous reports suggest that this catalytic loop configuration is stabilized by the highly conserved residues V809, P854, and E884 from the adjacent protease domain (**Fig. 4d** and **Supplementary Fig. 7**)^25,28^. Consistent with an important role in protease activation, mutating these residues to alanine severely impairs human LONP1 proteolytic activity, while only minimally affecting ATP hydrolysis (**Fig. 4e-f**).

To confirm and further characterize the dynamics of protease domain configurations induced by bortezomib, we performed hydrogen-deuterium exchange mass spectrometry (HDX-MS) in the presence and absence of bortezomib (**Fig. 4f** and **Supplementary Fig. 12**). Consistent with our structures of substrate-bound LONP1 and LONP1^Bz^, HDX-MS shows that the primary effect of bortezomib is to induce conformational remodeling of the protease domain. Notably, peptides comprising residues within the two helices flanking the catalytic S855-containing loop and the loop itself (peptides 803-820, 836-859, and 885-909) show significant stabilization upon addition of bortezomib (**Fig. 4g** and **Supplementary Fig. 12**), reflecting the importance of these residues in maintaining the active form of proteolytic active site. These results, along with our structures of substrate-bound LONP1 and LONP1^Bz^, support a model whereby engagement of substrate at the protease active site promotes conformational remodeling that includes inter-subunit stabilization of the autoinhibitory loop region to promote proteolytic activation of human LONP1.

## Discussion

Here, we determine structures for human LONP1 in three different states: substrate-free, substrate-bound, and both substrate- and bortezomib-bound. These structures allowed us to establish a molecular framework to define the proteolytic activation of human LONP1. Surprisingly, despite being highly conserved, our structures identify key differences between human and bacterial Lon that likely reflect evolutionary adaptations important for mitochondrial proteostasis regulation. We show that human LONP1 integrates a second pore loop residue that allows increased interactions with the translocating peptide. These increased interactions likely provide the human protease enhanced ‘grip’ on substrates, potentially improving its ability to degrade damaged or nonnative proteins within the mitochondrial environment, such as oxidatively-modified proteins previously shown to be LONP1 substrates^10,12^.

Further, we show that, unlike bacterial Lon, substrate engagement by the AAA+ domains of human LONP1 is not sufficient to promote conformational rearrangements within the protease domain required for relief of protease auto-inhibition. Instead, formation of a substrate-binding cleft at the protease active site appears to be linked to substrate interaction with the active site, suggesting a second level of regulation for human LONP1 that could function to prevent the aberrant degradation of proteins important for mitochondrial function (**Fig. 5**). Indeed, given the diversity of substrates processed by human LONP1 within the mitochondrial matrix, substrate-induced activation of individual protease active sites would provide a powerfully effective means by which to increase substrate selectivity and prevent aberrant degradation of mistargeted substrates. It is thus possible that all six protease active sites of human LONP1 are rarely, if ever, simultaneously activated for proteolytic activity. Such a system would endow human LONP1 with a level of proteolytic plasticity to enable enzymatic activity to be specifically tuned to the types and quantities of substrates present within the matrix, which continuously fluctuate in response to cellular queues.

**Figure 5.**
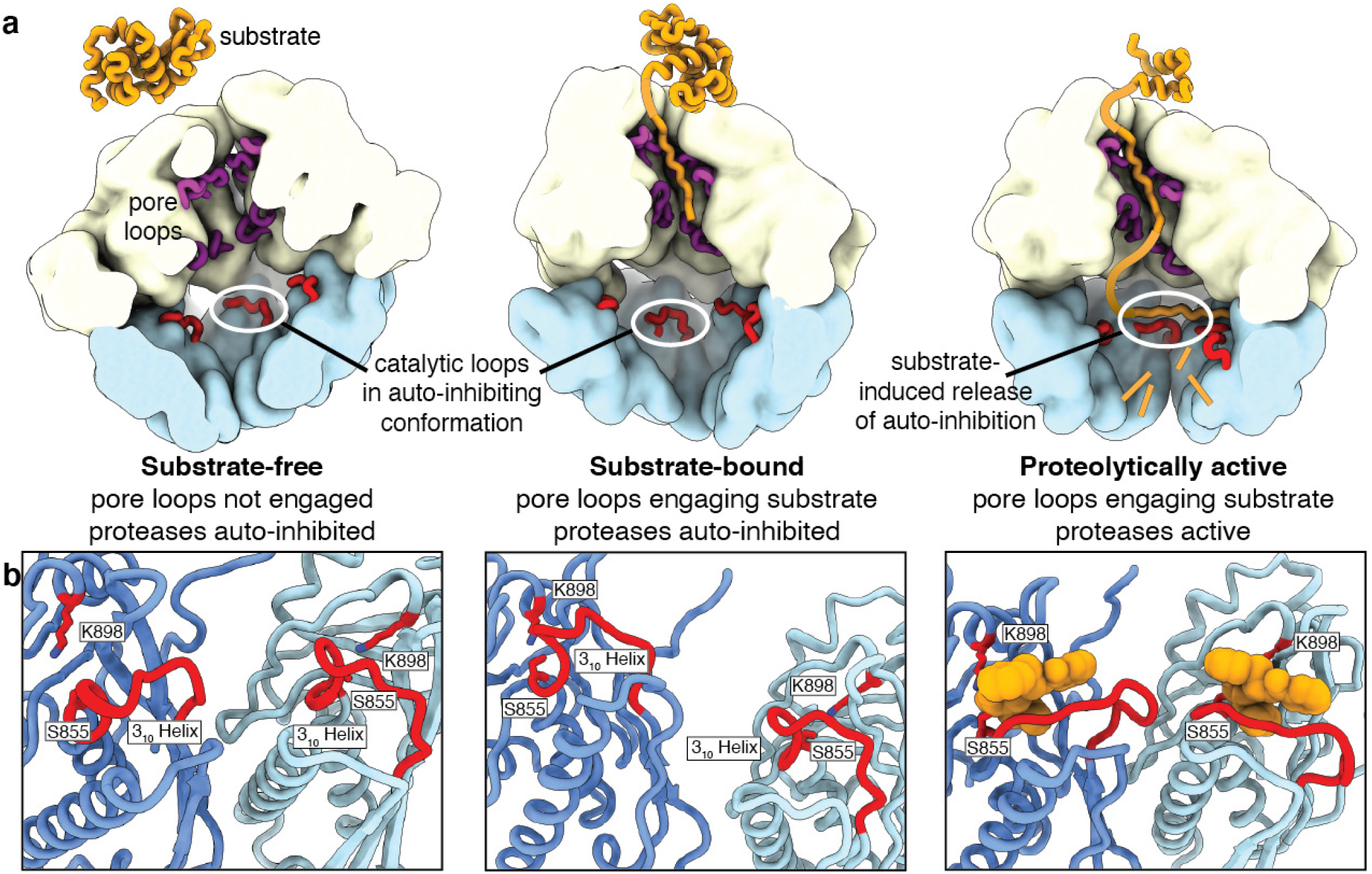
Model for protease activation in LONP1. **a.** Substrate is engaged by the open state conformer (left), causing a transition to a translocation-competent organization of the ATPases (middle). Despite substrate engagement by the pore loops (purple) in the central channel, the protease active sites remain in an auto-inhibited conformation. This auto-inhibition is only relieved upon interaction with substrate (right), at which point the catalytic loop (red) extends away from the catalytic residues to enable substrate cleavage. **b.** Close-up views of the proteolytic active sites of substrate-free, substrate-bound, and proteolytically active LONP1 enzymes. In the auto-inhibited substrate-free and substrate-bound states, a 310 helix sterically hinders substrates from entering the proteolytic active site for degradation in addition to preventing the formation of the catalytic dyad between S855 and K898. However, in the presence of substrate (represented by bortezomib in orange in the right panel) in the proteolytic active site, auto-inhibition releases as the catalytic serine-containing loop extends towards the neighboring subunit, the catalytic dyad is formed, and the protease domains activate for proteolytic cleavage.

Lastly, our structures also provide a structural framework for defining the pathologic and potentially therapeutic implications of LONP1 in human disease. For example, mapping known CODAS syndrome-associated mutations of human *LONP1* onto our structure of proteolytically active LONP1^Bz^ highlights their position at interfaces between ATPase domains (**Supplementary Fig. 13**). This suggests that these mutations likely impact the tight subunit coordination required for ATP hydrolysis and substrate translocation. Consistent with this, CODAS mutations were previously shown to reduce LONP1 ATPase and proteolytic activities^17^. Given that the diversity of CODAS mutations, as well as the unique combination of mutations present in heterozygous individuals, result in a wide array of phenotypes in affected individuals, our atomic models of human LONP1 provide the structural basis for future analyses of CODAS patient mutations required to understand the diverse etiology of this disease. Further, our structures provide a molecular framework to facilitate the development of small molecule strategies to modulate LONP1 activity to mitigate pathologic mitochondrial dysfunctions associated with imbalances in LONP1 proteolysis^14,42^.

## Materials and Methods

### Protein expression and purification

A gene encoding an N-terminally His_6_-tagged human LONP1 with methionine 115 (the N-terminal residue of mature mitochondrial-matrix localized LONP1) as its first residue was inserted into a pET20b *E. coli* expression vector. The pET20b-*His*_6_*-LONP1* plasmid was transformed into the Rosetta 2(DE3)pLysS *E. coli* strain for recombinant protein expression. Cells were grown in 3% tryptone, 2% yeast extract, and 1% MOPS pH 7.2 with 100 μg/ml ampicillin and 25 μg/ml chloramphenicol and cultured at 37°C shaking at 220 rpm to an optical density (OD_600_) of 0.8. Protein overexpression was induced with 1 mM IPTG for 16 hours at 16°C. Cells were harvested by centrifugation at 5,000 x g, resuspended in buffer A (0.2M NaCl, 25mM Tris pH 7.5, 20% glycerol, and 2mM β-mercaptoethanol), flash-frozen and then stored at −80°C for later use. Upon thawing, cells were lysed by sonication. Cell lysate was cleared by centrifugation at 15,000 rpm and then added to TALON Cobalt resin equilibrated in Buffer A and loaded onto a gravity-flow column to allow His_6_-LONP1 binding. The column was washed with buffer A containing 20 mM imidazole to remove unbound proteins. Bound proteins were eluted in 1.2 column volumes of buffer A containing 300 mM imidazole. Eluted proteins were concentrated and loaded onto a Superose 6 size exclusion column equilibrated with buffer A. Fractions containing His_6_-LONP1 were pooled, concentrated, and flash-frozen for storage at −80°C. The pET20b-*His*_6_*-LONP1* plasmid served as a template for generating *LONP1* mutants using the quick-change site directed mutagenesis approach. Each LONP1 mutant protein was expressed and purified as described above.

### *In vitro* proteolysis assay

Purified human LONP1 (0.1 μM) was incubated for 15 minutes at 37°C in LONP1 activity buffer (50 mM Tris-HCl pH 8, 100 mM KCl, 10 mM MgCl_2_, 1 mM DTT, and 10% Glycerol) in the presence or absence of 2.5 mM ATP. FITC-Casein (0.8 μM; Sigma-Aldrich) was added to initiate the degradation reaction. An increase of fluorescence (excitation 485 nm, emission 535 nm) resulting from free FITC molecules was monitored using a TECAN plate reader. Three biological replicates were performed for each LONP1 mutant and the data were fit to a straight line from which the slope was extracted to calculate the rate of substrate degradation. GraphPad Prism software was used for data analysis. Mean and standard error of mean (SEM) was calculated by performing column statistics. ANOVA was used to calculate p-value, as indicated.

### *In vitro* ATP hydrolysis assay

*In vitro* ATP hydrolysis was carried out in LONP1 activity buffer (50 mM Tris-HCl pH 8, 100 mM KCl, 10 mM MgCl_2_, 10% Glycerol, and 1 mM DTT) in the presence or absence of 2.5 mM ATP. Casein (9.7 μM) was added to the reaction mixture when measuring substrate-induced ATPase activity. Reaction components and purified LONP1 (0.1 μM) were incubated separately at 37^*c*^*irc*C for five minutes. LONP1 was added to initiate ATP hydrolysis and the amount of free inorganic phosphate at each timepoint was measured by adding Malachite Green Working Reagent (Sigma-Aldrich) and incubating components for 30 minutes at room temperature for color development before measuring absorbance at 620 nm (OD_620_). Three biological replicates were performed for each LONP1 mutant in the presence and absence of substrate. Data were fit to a straight line and the slope were extracted to calculate ATP hydrolysis rates. GraphPad Prism software was used for data analysis. Mean and standard error of mean (SEM) was calculated by performing column statistics. ANOVA was used to calculate p-value, as indicated.

### Hydrogen-deuterium exchange (HDX) detected by mass spectrometry (MS)

Differential HDX-MS experiments were conducted as previously described with a few modifications^43^. Peptides were identified using tandem MS (MS/MS) with an Orbitrap mass spectrometer (Q Exactive, ThermoFisher). Product ion spectra were acquired in data-dependent mode with the top five most abundant ions selected for the product ion analysis per scan event. The MS/MS data files were submitted to Mascot (Matrix Science) for peptide identification. Peptides included in the HDX analysis peptide set had a MASCOT score greater than 20 and the MS/MS spectra were verified by manual inspection. The MASCOT search was repeated against a decoy (reverse) sequence and ambiguous identifications were ruled out and not included in the HDX peptide set.

For HDX-MS analysis, LONP1 (10 μM) was incubated with the respective ligands at a 1:10 protein-to-ligand molar ratio for 1 h at room temperature. Next, 5 μl of sample was diluted into 20 μl D_2_O buffer (50 mM Tris-HCl, pH 8; 75 mM KCl; 10 mM MgCl2) and incubated for various time points (0, 10, 60, 300, and 900 s) at 4°C. The deuterium exchange was then slowed by mixing with 25 μl of cold (4°C) 0.1M Sodium Phosphate, 50mM TCEP. Quenched samples were immediately injected into the HDX platform. Upon injection, samples were passed through an immobilized pepsin column (2mm × 2cm) at 200 μl min^−1^ and the digested peptides were captured on a 2mm × 1cm C_8_ trap column (Agilent) and desalted. Peptides were separated across a 2.1mm × 5cm C_18_ column (1.9 μl Hypersil Gold, ThermoFisher) with a linear gradient of 4% - 40% CH_3_CN and 0.3% formic acid, over 5 min. Sample handling, protein digestion and peptide separation were conducted at 4°C. Mass spectrometric data were acquired using an Orbitrap mass spectrometer (Exactive, ThermoFisher). HDX analyses were performed in triplicate, with single preparations of each protein ligand complex. The intensity weighted mean m/z centroid value of each peptide envelope was calculated and subsequently converted into a percentage of deuterium incorporation. This is accomplished determining the observed averages of the undeuterated and fully deuterated spectra and using the conventional formula described elsewhere^44^. Statistical significance for the differential HDX data is determined by an unpaired t-test for each time point, a procedure that is integrated into the HDX Workbench software^45^. Corrections for back-exchange were made on the basis of an estimated 70% deuterium recovery, and accounting for the known 80% deuterium content of the deuterium exchange buffer.

For data rendering, HDX data from all overlapping peptides were consolidated to individual amino acid values using a residue averaging approach. Briefly, for each residue, the deuterium incorporation values and peptide lengths from all overlapping peptides were assembled. A weighting function was applied in which shorter peptides were weighted more heavily and longer peptides were weighted less. Each of the weighted deuterium incorporation values were then averaged to produce a single value for each amino acid. The initial two residues of each peptide, as well as prolines, were omitted from the calculations. This approach is similar to that previously described^46^. HDX analyses were performed in triplicate, with single preparations of each purified protein/complex. Statistical significance for the differential HDX data is determined by t test for each time point, and is integrated into the HDX Workbench software^45^.

### Sample preparation for electron microscopy

Wildtype human LONP1 was diluted to a concentration of 2.5 mg/ml in 50 mM Tris pH 8, 75 mM KCl, 10 mM MgCl_2_, 1 mM TCEP, and 1 mM ATPγS. 4 μl of the sample were applied onto 300 mesh R1.2/1.3 UltrAuFoil Holey Gold Films (Quantifoil) that were plasma cleaned prior to sample application for 7 seconds using a Solarus plasma cleaner (Gatan, Inc.) with a 75% nitrogen, 25% oxygen atmosphere at 15W. Excess sample was blotted away for 4 s using Whatman No. 1 filter paper and vitrified by plunge freezing into a liquid ethane slurry cooled by liquid nitrogen using a manual plunger in a 4° C cold room whose humidity was raised to 95% using a humidifier. For bortezomib-bound LONP1, LONP1 was similarly diluted to a concentration of 2.5 mg/ml in 50 mM Tris pH 8, 75 mM KCl, 10 mM MgCl_2_, 1 mM TCEP, and 1 mM ATPγS with the addition of 10-fold molar excess bortezomib. Samples were prepared for cryo-EM analyses using the same procedures used for the wildtype sample.

### Electron microscopy data acquisition

For wildtype human LONP1, cryo-EM data were collected on a Thermo-Fisher Talos Arctica transmission electron microscope operating at 200 keV using parallel illumination conditions^47^. Micrographs were acquired using a Gatan K2 Summit direct electron detector, operated in electron counting mode applying a total electron exposure of 50 e^−^/Å^2^ as a 114-frame dose-fractionated movie during a 11.4s exposure (**Supplementary Fig. 1a**). The Leginon data collection software^48^ was used to collect 2914 micrographs at 36,000x nominal magnification (1.15 Å /pixel at the specimen level) with a nominal defocus set to −1.5 μm. Variation from the nominally set defocus due to a 5% tilt in the stage gave rise to a defocus range (this tilt was not intentional and required service to correct). Stage movement was used to target the center of sixteen 1.2 μm holes for focusing, and an image shift was used to acquire high magnification images in the center of each of the sixteen targeted holes.

Similarly for bortezomib-bound LONP1, cryo-EM data were collected on a Thermo-Fisher Talos Arctica transmission electron microscope operating at 200 keV using parallel illumination conditions^47^. Micrographs were acquired using a Gatan K2 Summit direct electron detector, operated in electron counting mode applying a total electron exposure of 50 e^−^/Å^2^ as a 52-frame dose-fractionated movie during a 10.4 s exposure (**Supplementary Fig. 1a**). The Leginon data collection software^48^ was used to collect 4776 micrographs at 36,000x nominal magnification (1.15 Å /pixel at the specimen level) with a nominal defocus of −1.2 μm (a single value due to the tilted stage). Images for bortezomib-bound LONP1 were collected using a similar strategy for wildtype LONP1. Micrograph frames were aligned using MotionCor2^49^ and CTF parameters were estimated with CTFFind4^50^ in real-time during data acquisition to monitor image quality using the Appion image processing environment^51^.

### Image processing for the wildtype human LONP1 + ATPγS

A small dataset of 40,000 particles were picked using a difference of gaussians picker^52^, which was used for reference-free 2D classification in Appion49. Representative views were selected as templates for template-based particle picking using FindEM^53^, which yielded 940,396 putative particle selections. All subsequent processing was performed in RELION 3.1^54^. Particle coordinates were extracted at 3.45 Å /pixel from the motion-corrected/dose-weighted micrographs with a box size of 80 pixels, and 2D classified into 200 classes. Based on this 2D analysis, 564,930 particles belonging to classes displaying details corresponding to secondary structural elements were selected for further processing in 3D (examples shown in **Supplementary Fig. 1b**). A low-resolution negative stain reconstruction of human LONP1 was used as an initial model for 3D classification of the particles (3 classes, T=4, 25 iterations). One 3D class containing 52% of the particles resembled an open lockwasher (hereafter referred to as substrate-free), while the other two classes resembled the previously determined substrate-bound Lon complex^28^ (**Supplementary Fig. 1b**). The alignment parameters were used to extract 293,886 and 271,044 centered, full-size particles corresponding to the substrate-free and substrate-bound complexes, respectively (1.15 Å /pixel, box size 288 pixels). The substrate-free and substrate-bound 3D classes were scaled to the appropriate pixel size and used as initial models for 3D auto-refinement of these particles, resulting in reconstructions at reported resolutions of 4.9 and 4.7 Å according to FSC of half maps at 0.143, respectively. The image shifts imposed to acquire each group of 16 exposures during data collection were used to generate 16 optics groups for CTF refinement^55^ (per-particle defocus, per-micrograph astigmatism, and beam-tilt estimation). CTF refinement followed by 3D auto-refinement using a soft-edged 3D mask was repeated three times, ultimately resulting in reconstructions with reported resolutions of 3.4 Å and 3.2 Å for the substrate-free and substrate-bound complexes, respectively (**Supplementary Fig. 1b**).

Notably, the density for one of the six subunits in the substrate-free reconstruction was almost non-existent, which prompted us to perform a 3D classification without alignment on the particles contributing this reconstruction (3 classes, T=15, 50 iterations). One of the three classes contained stronger density corresponding to the sixth subunit, and the subset of 45,300 particles contributing to this reconstruction were selected for 3D auto-refinement. The resulting six-subunit structure was determined to have a resolution of 3.6 Å, although the sixth subunit remained insufficiently resolved for *ab initio* modeling (colored blue in **Supplementary Fig. 1b**).

3D classification without alignment was used to identify the particles containing the highest resolution structural data contained in both the substrate-free and substrate-bound particle stacks. 3D masks encompassing the five subunits of the substrate-free complex, and all six subunits for the substrate-bound complex were used for these classifications (3 classes, T=15, 50 iterations). 83,162 particles from the two highest-resolution classes of the substrate-free complex, and 38,130 particles from the highest resolution class of the substrate-bound complex were selected for a final masked 3D auto-refinement using local angular and translational searches (0.9 degrees, 2 pixel search range with a 0.5 pixel step). The resulting reconstructions showed a marginal qualitative improvement in density, although the reported resolutions were unchanged at 3.4 and 3.2 Å for the substrate-free and substrate-bound, respectively (Supplementary Fig. 1c).

### Image processing for the wildtype human LONP1 + ATPγS + bortezomib

Micrograph frames were aligned using MotionCor2^49^ and CTF parameters were estimated with gCTF^56^ using the RELION 3.1 processing environment^54^. 2D classes representing orthogonal views of the human LONP1 complex were used for template-based particle selection in RELION. 2,736,565 particles were selected from micrographs and extracted using a box size of 256 pixels. The particle stack was binned by a factor of 4 for reference-free 2D alignment and classification (**Supplementary Fig. 9a**), and 1,539,141 particles were selected for initial processing. A low-resolution negative stain reconstruction of human LONP1 was used as an initial model for 3D refinement of particles. These particles refined to a reported resolution of 9.5 Å as estimated by FSC at a cutoff of 0.143. The particles from this reconstruction were then sorted by classification with alignment into five classes. The particles from this reconstruction were then sorted by classification with alignment into five classes, two of which, accounting for 92% of particles, displayed high-resolution features that did not contain anisotropic resolution artifacts. The 1,418,681 particles from these classes were merged and refined to 5.0 Å resolution. 3D classification without alignment was performed to increase resolution of the reconstruction. 1,142,531 particles were selected for further refinement. The X and Y image shifts applied during data acquisition were used to group images for beam tilt estimation. Refining with local defocus and beam tilt estimation within CTF Refinement in RELION 3.1 improved the overall reported resolution of our reconstruction of the substrate-bound human LONP1 to 4.1 Å by FSC at 0.143. Several more rounds of 3D classification without alignment were performed to continue increasing resolution of the reconstruction. 532,298 particles were selected for the final 3D refinement and Bayesian Polishing, resulting in a final reconstruction of bortezomib-bound human LONP1 with an overall reported resolution of 3.7 Å by FSC at 0.143 (**Supplementary Fig. 9c**). The cryo-EM density of the two seam subunits was poorly resolved in this reconstruction, so a soft-edged 3D mask was generated to encompass the step subunits for focused refinement. Focused refinement improved the resolution in this region by 0.4 Å as shown by local resolution estimation in RELION 3.1 (**Supplementary Fig. 9d**).

### Atomic model building and refinement

Model building and refinement were similarly performed for the substrate-bound and bortezomib-bound LONP1 structures. A homology model of human LONP1 was generated using a previously resolved cryo-EM structure of bacterial LONP1 as a starting model using SWISS-MODEL^57^. This initial model was split into ATPase and protease domains and rigid body docked into the density of each of the subunits using UCSF Chimera^58^. Real-space refinement of the docked structures and *ab initio* model building were performed in COOT^59^. The flexible linker regions were modeled *ab initio* and a twelve-amino acid poly-alanine peptide as well as ATP, ADP, magnesium cofactor, and bortezomib molecules were built into the density corresponding to substrate, nucleotide, and drug, respectively. Further refinement of the full hexameric atomic model was performed using one round each of morphing and simulated annealing in addition to five real-space refinement macrocycles with atomic displacement parameters, secondary structure restraints, local grid searches, non-crystallographic symmetry, Ramachandran restraints, and global minimization in PHENIX^60^. One round of geometry minimization with Ramachandran and rotamer restraints was used to minimize clash scores, followed by a final round of real-space refinement in PHENIX.

To create an initial model for substrate-free LONP1, individual subunits from the substrate-bound human LONP1 atomic model were docked into the density of each of the five well-resolved subunits of substrate-free LONP1 using UCSF Chimera^56^. Real-space refinement of the docked structures and *ab initio* model building were performed in COOT^57^. Residues from the model in were trimmed to Cα unless there was clearly discernible EM density for the Cβ atom. Further refinement of the model was performed using one round each of morphing and simulated annealing in addition to five real-space refinement macrocycles with atomic displacement parameters, secondary structure restraints, local grid searches, non-crystallographic symmetry, Ramachandran restraints, and global minimization in PHENIX^60^. ADP molecules were built into the density corresponding to nucleotide into the five well-resolved subunits. One round of geometry minimization with Ramachandran and rotamer restraints was used to minimize clash scores followed by a final round of real-space refinement in PHENIX. Due to the low resolution of the sixth subunit (blue in **Supplementary Fig. 1b**) atomic coordinates for this subunit are not included in the deposited atomic model. However, since secondary structural elements were visible for this subunit in the reconstruction, a copy of subunit B from our five-subunit substrate-free model was docked into the density for the sixth subunit for interpretation and figure-making.

UCSF Chimera^58^ and ChimeraX^61^ were used to interpret the EM reconstructions and atomic models, as well as to generate figures.

## Data availability

Maps for the substrate-free, substrate-bound and LONP1^Bz^ complexes were deposited in the Electron Microscopy Data Bank under accession IDs EMD-23019, EMD-23020, and EMD-23013, respectively. Corresponding atomic models were deposited in the Protein Data Bank under accession IDs 7KSL, 7KSM, and 7KRZ, respectively.

## Acknowledgements

We thank J.C. Ducom at Scripps Research High Performance Computing and C. Bowman at Scripps Research for computational support, as well as B. Anderson at the Scripps Research Electron Microscopy Facility for microscopy support. We thank Nigel Moriarty for assistance in refinement of the LONP1 bortezomib-bound structure. Further we thank Carolyn Suzuki (Rutgers) and members of the Lander and Wiseman labs at Scripps Research for insightful and helpful discussions related to this work. M.S. is supported by the National Science Foundation Graduate Research Fellowship Program. G.C.L. is supported by the National Institutes of Health (NIH) AG067594 and an Amgen Young Investigator Award. R.L.W. is supported by the NIH NS095892. G.C.L. and R.L.W. are supported by NIH AG061697. Computational analyses of EM data were performed using shared instrumentation funded by NIH S10OD021634 to G.C.L.

## Supplementary Materials

**Supplementary Figure 1.**
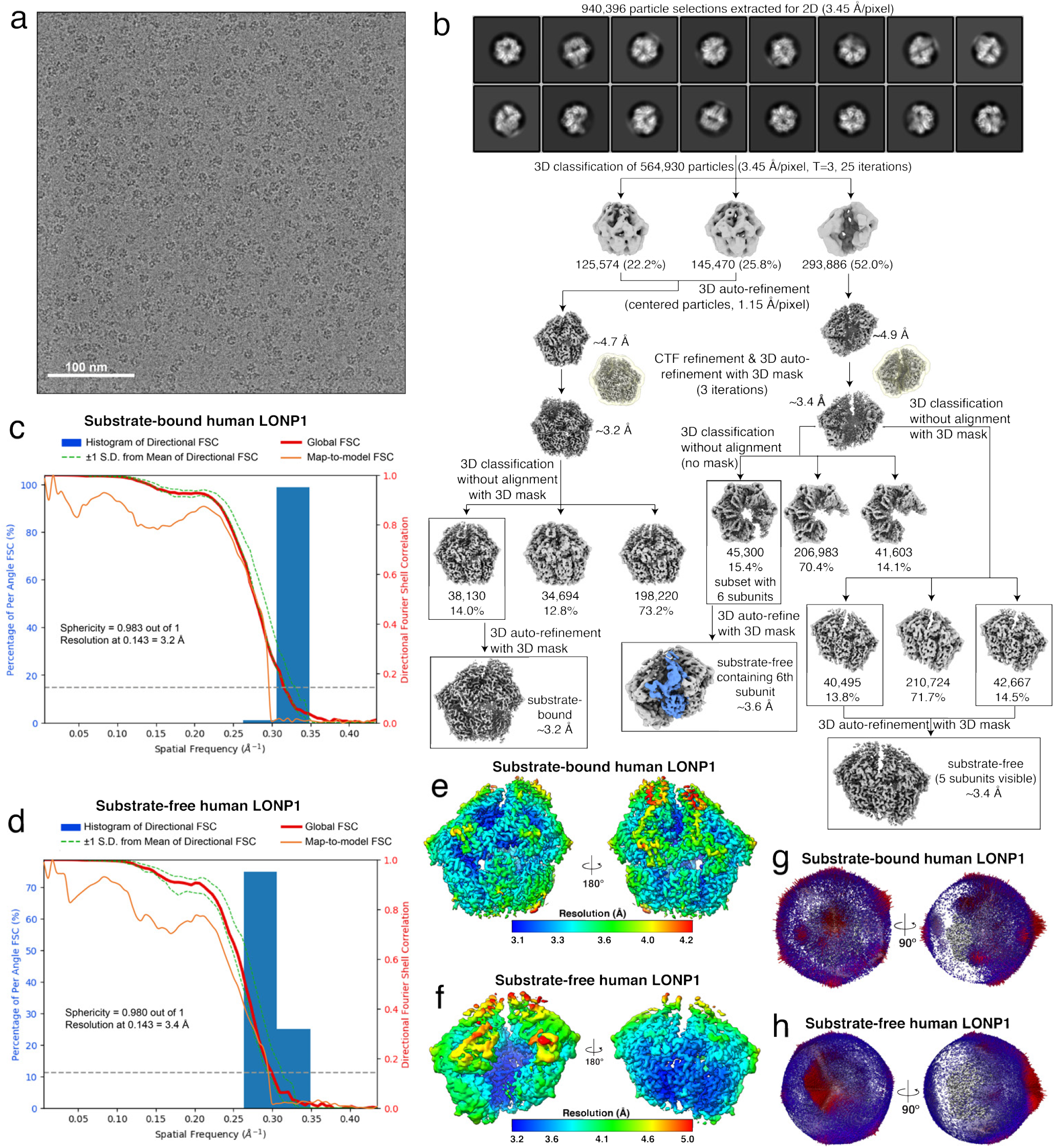
Cryo-EM structure determination of the human LONP1 substrate-free and substrate-bound complexes. **a.** Representative micrograph from cryo-EM data collection. **b.** Cryo-EM data processing scheme followed using RELION 3.1 software^54^ to obtain the final 3D reconstructions of substrate-bound and substrate-free human LONP1, including classification revealing a subset of substrate-free particles where the sixth subunit is visible (colored blue). The final maps were used for atomic model building and refinement. **c-d.** 3-Dimensional Fourier Shell Correlation (3DFSC)62 of the final reconstruction of substrate-bound (**c**) and substrate-free (**d**) human LONP1 reporting a global resolution of 3.2 Å and 3.4 Å at FSC=0.143, respectively. Both reconstructions showed a high level of sphericity, as calculated by the 3DFSC (0.983 out of 1 for the substrate-bound and 0.980 out of 1 for the substrate-free). **e-f.** Final substrate-bound (**e**) and substrate-free (**f**) reconstructions filtered and colored by local resolution, as calculated using RELION. The final EM densities are resolved to a range of resolutions. The substate-bound complex is mostly resolved to 3.1 Å resolution at the core of the complex, but > 4.0 Å in more flexible regions, such as the ‘seam’ subunit. In the substrate-free reconstruction, the protease domains of the central subunits are mostly resolved to 3.2 Å resolution, while the peripheral subunits and most of the ATPase domains are resolved to > 4.0 Å. **g-h.** Euler angle distribution plots of the 38,130 and 83,162 particles used in the final reconstruction of the substrate-bound and substrate-free structures, shown from two orthogonal directions.

**Supplementary Figure 2.**
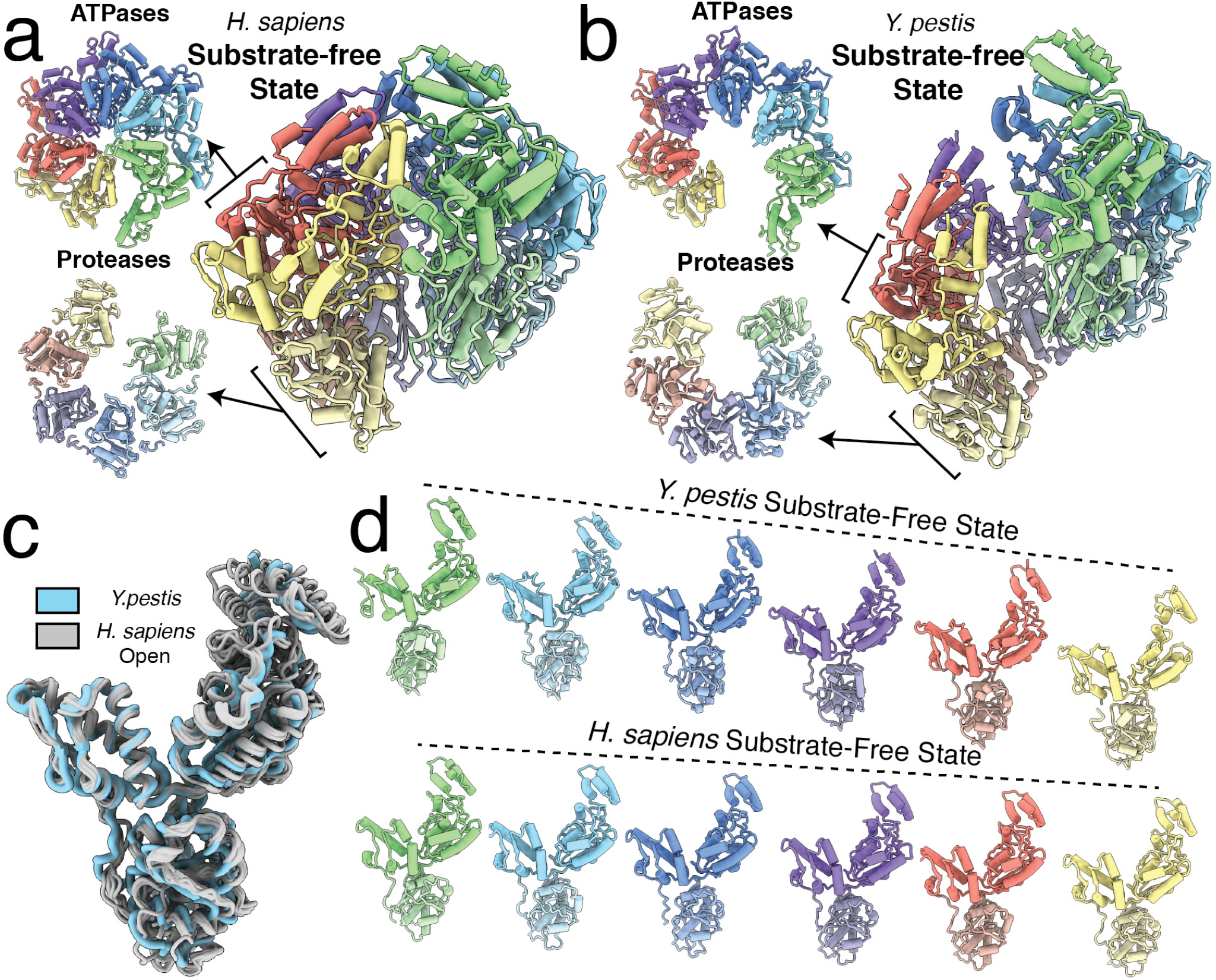
Comparison of substrate-free Lon protease structures from *Y. pestis* and *H. sapiens*. *a.*Axial (left) and lateral (right) views of the substrate-free human LONP1 atomic model (**a**) next to the *Y. pestis* Lon substrate-free atomic model^28^ (**b**). Axial views of the ATPase and protease domain rings are shown from the exterior of each ring. Subunits are colored as in **Fig. 1**. **c.** Individual subunits from the substrate-free human LONP1 structure (grey) were aligned using a secondary structure-based alignment and subsequently aligned to a subunit from the substrate-free *Y. pestis* structure (light blue). These alignments reveal structural conservation between substrate-free states Lon homologs. **d.** Individual protomers of substrate-free bacterial Lon (top) and substrate-free human LONP1 (bottom) in the intermediate state lined alongside one another relative to the protease domain produced by orienting all the protease domains to a common view. The subunits of the substrate-free *Y. pestis* structure possess a steeper left-handed helical pitch than in the human substrate-free state.

**Supplementary Figure 3.**
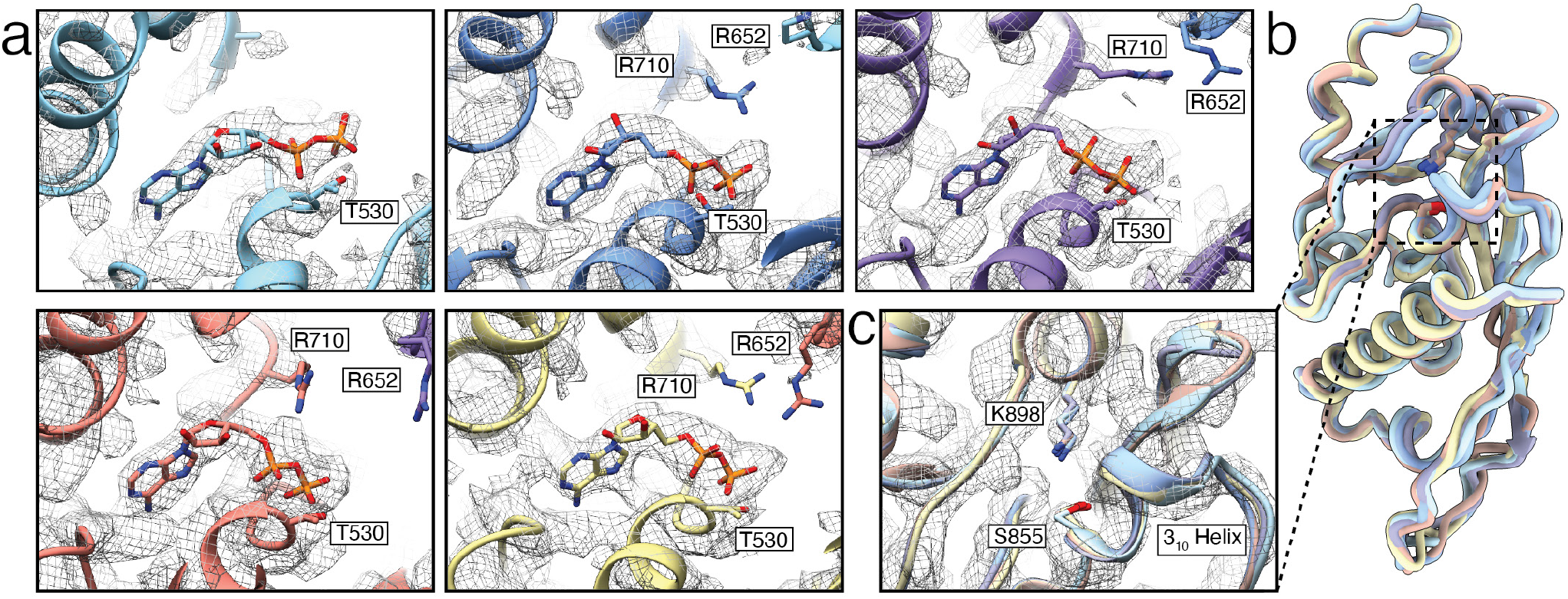
Structure of substrate-free LONP1 represents a fully ADP-bound, inactivated conformation. The nucleotide binding pockets of all five subunits of substrate-free LONP1 arranged from highest (light blue) to lowest (yellow) subunits, with the cryo-EM density in this region shown as an isosurface mesh contoured at sigma = 3.6. All subunits possess density for nucleotide that is unambiguously consistent with an ADP molecule. **b.** Alignment of all five protease domains of the substrate-free state of LONP1 show that the protease domain adopts a consistently inactivated conformation where a 310 helix blocks entrance of substrates into the proteolytic active site, highlighted by a hatched box. **c.** A zoomed-in view of the proteolytic active sites of all six subunits exhibiting the auto-inhibited conformation of LONP1.

**Supplementary Figure 4.**
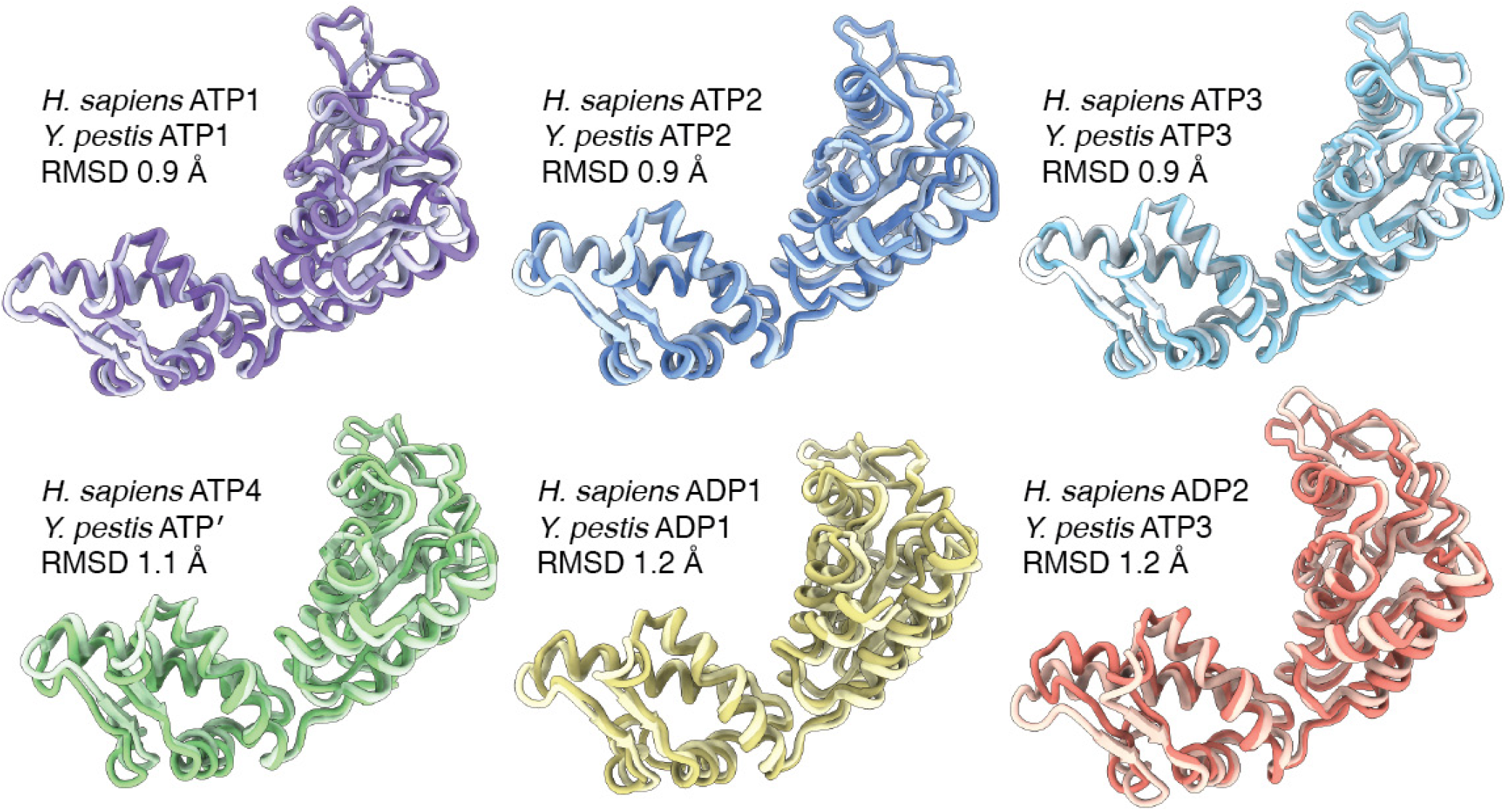
ATPase domains of human LONP1 have a nearly identical organization as *Y. pestis* Lon. The AAA+ of the substrate-bound LONP1 conformer are aligned to the corresponding AAA+ domains from substrate-bound *Y. pestis* Lon28. All six subunits of the ATPase overlay with RMSDs <1.2 Å.

**Supplementary Figure 5.**
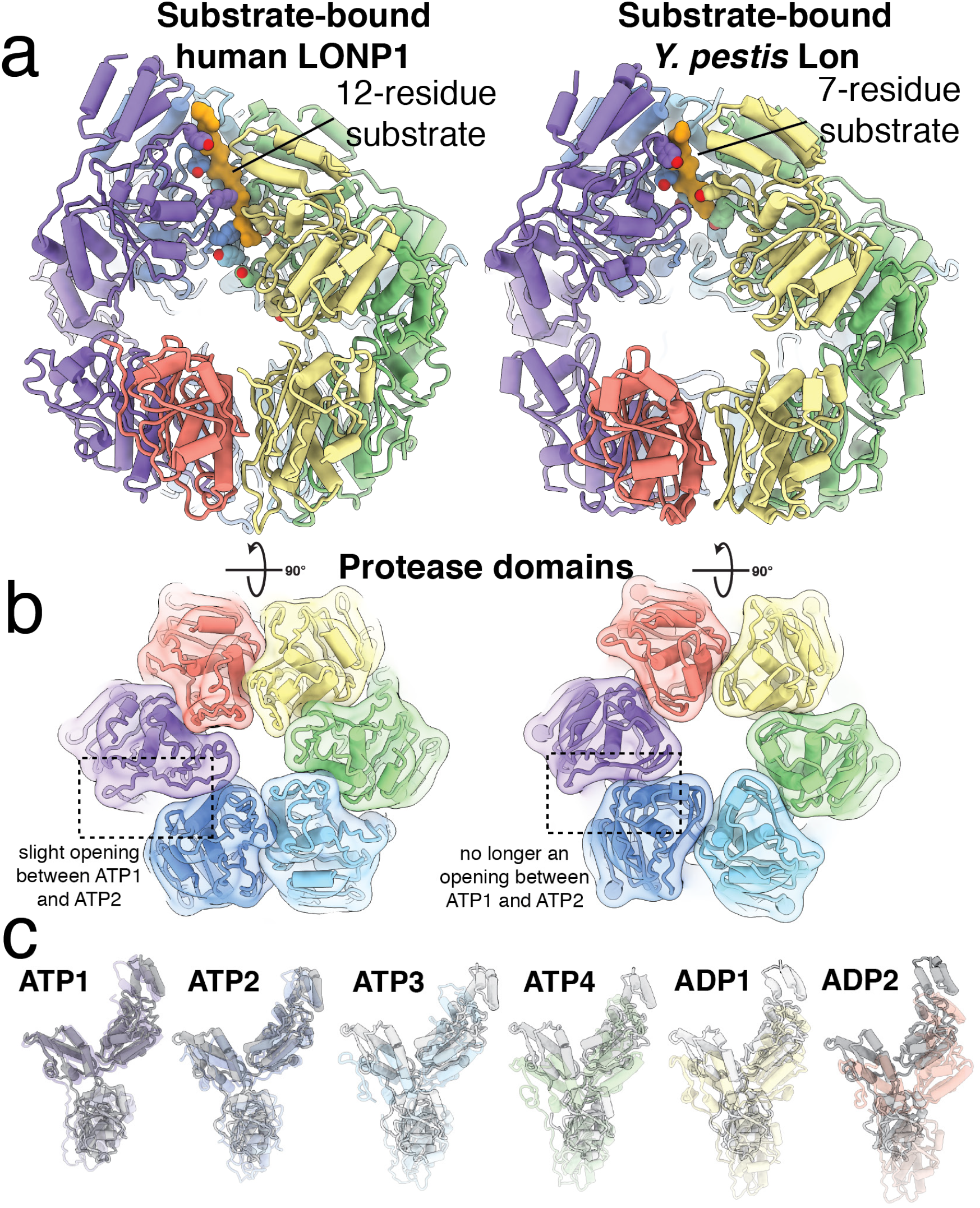
Comparison of substrate-bound Lon protease structures from *Y. pestis* and *H. sapiens*. **a.** Cutaway views of the substrate-bound human LONP1 and substrate-bound *Y. pestis* Lon atomic models28. Subunits are colored as in **Fig. 1.** Notably, human LONP1 has a 12-residue substrate trapped in its central channel while *Y. pestis* contains 7 residues, likely due to additional substrate interactions evolved in human LONP1, namely, a tyrosine pore-loop 2. **b.** Axial views of the protease rings emphasize the asymmetry of the substrate-bound LONP1 (left), which has a slight opening between the protease subunits of ATP1 and ATP2, compared to the six-fold symmetric organization of the substrate-bound *Y. pestis* Lon (right). **c.** Individual protomers of substrate-bound bacterial Lon aligned side-by-side relative to the protease domain produced by orienting all the proteases to a common view. Subunits from substrate-bound human LONP1 are shown colored as in (**a**) and shown with transparency on top of the subunits of bacterial Lon (colored gray). While the spiraling ATPases sit atop a planar proteolytic ring in bacterial Lon, both AAA+ and protease domains in substrate-bound human LONP1 are asymmetric.

**Supplementary Figure 6.**
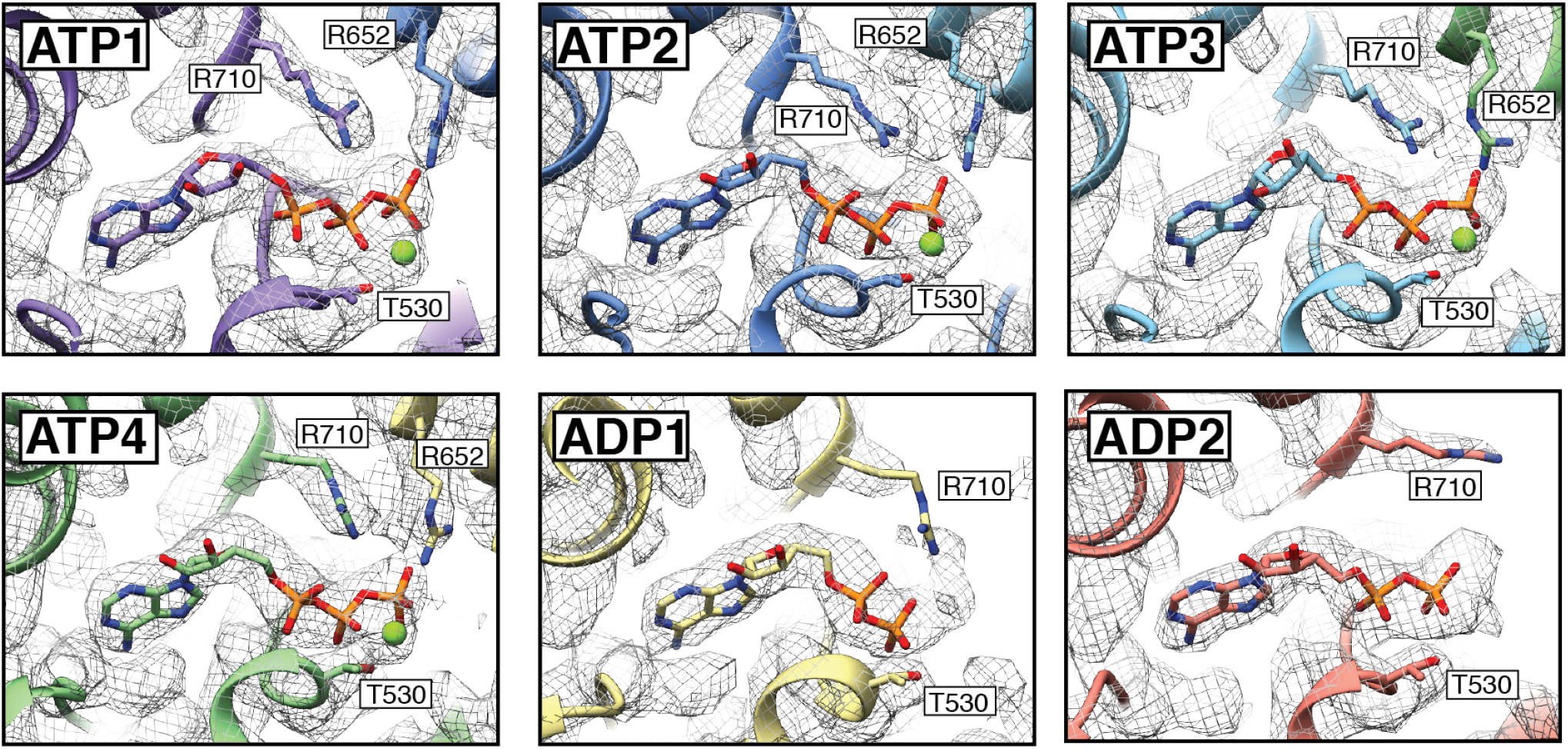
Substrate-bound human LONP1 shows nucleotide densities in the nucleotide-binding pocket. The cryo-EM density, shown as an isosurface mesh contoured at sigma=4.0, in the vicinity of the nucleotide binding pocket was of sufficient quality to assign the nucleotide state of each of the subunits. ATP1, ATP2, ATP3, and ATP4 subunits correspond to the ATPγS used to determine this structure coordinated by a magnesium cofactor. The nucleotide density in the ADP1 and ADP2 subunits likely corresponded to ADP molecules.

**Supplementary Figure 7.**
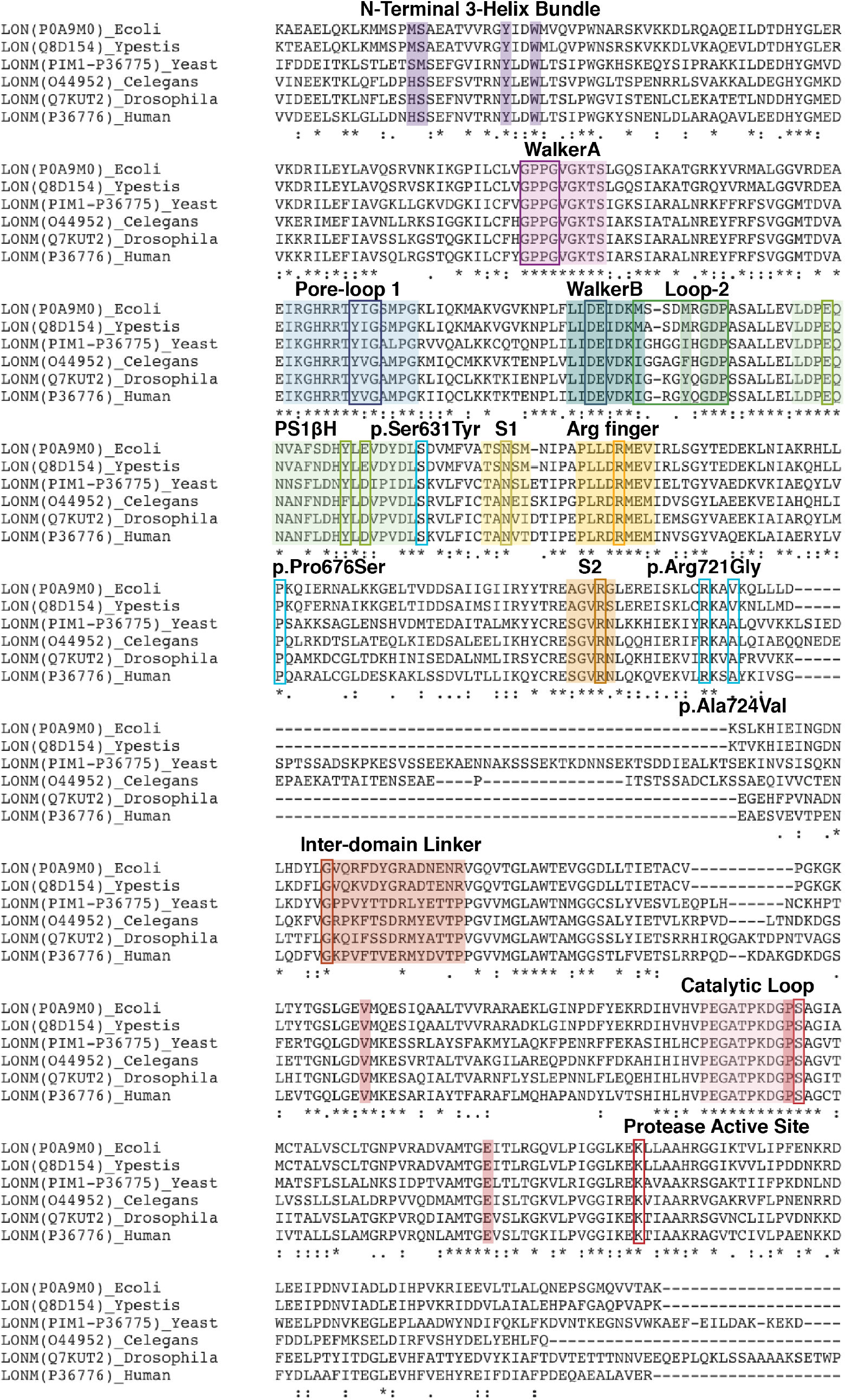
Sequence alignment of Lon protease homologs. ClustalW^63^ alignment of the Uniprot sequences of Lon homologs, including cytosolic Lon from *E. coli* and *Y. pestis*, mitochondrial Lon from yeast (Pim1), *C. elegans*, *Drosophila melanogaster*, and humans. This alignment suggests conservation across Lon proteins. Notably, we found several key residues to be strictly conserved in all sequences studied: the N-terminal 3-Helix Bundle (purple boxes around conserved residues studied in *Y. pestis*), P-loop (magenta), pore-loop 1 aromatic-hydrophobic residues (blue), the Walker B motif (teal), Loop-2 (green), pre-sensor 1 beta hairpin insertion (light green), sensor-1 (yellow), a trans-acting arginine finger (orange-yellow box), a cis-acting arginine finger present in sensor-2 (orange), an inter-domain linker containing a glycine residue at its N-terminus (red-orange), and a catalytic loop that is folded into a 3_10_ helix in the proteolytic active site in auto-inhibited Lon and extended in the activated conformation (salmon). Conserved residues involved in stabilizing the activated configuration of the protease domain are highlighted using filled red boxes (V809, P854, and E882). Mutations associated with CODAS syndrome are highlighted using cyan boxes: p.Ser631Tyr, p.Pro676Ser, p.Arg721Gly, and p.Ala724Val. Residues mutated in CODAS syndrome exhibit conservation across Lon homologs, suggesting their importance for Lon protease activity.

**Supplementary Figure 8.**
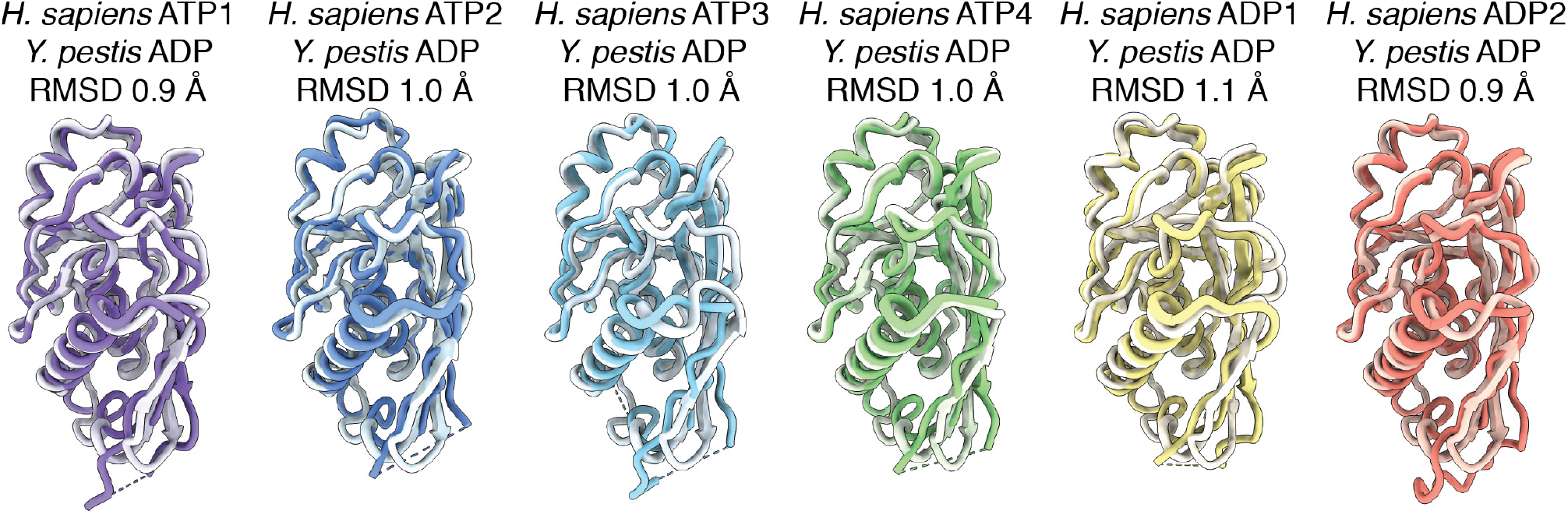
Protease domains in the substrate-bound LONP1 structure are auto-inhibited. The protease domains of substrate-bound human LONP1 aligned to their corresponding subunits from the substrate-free *Y. pestis* Lon^28^. All six subunits of the protease domain are in similar conformations, with RMSDs between 0.9-1.1 Å. Despite having substrate trapped in its central channel, the protease domain of semi-closed LONP1 is auto-inhibited by the catalytic loop forming a 3_10_ helix in the active site, similar to substrate-free forms of bacterial and human Lon.

**Supplementary Figure 9.**
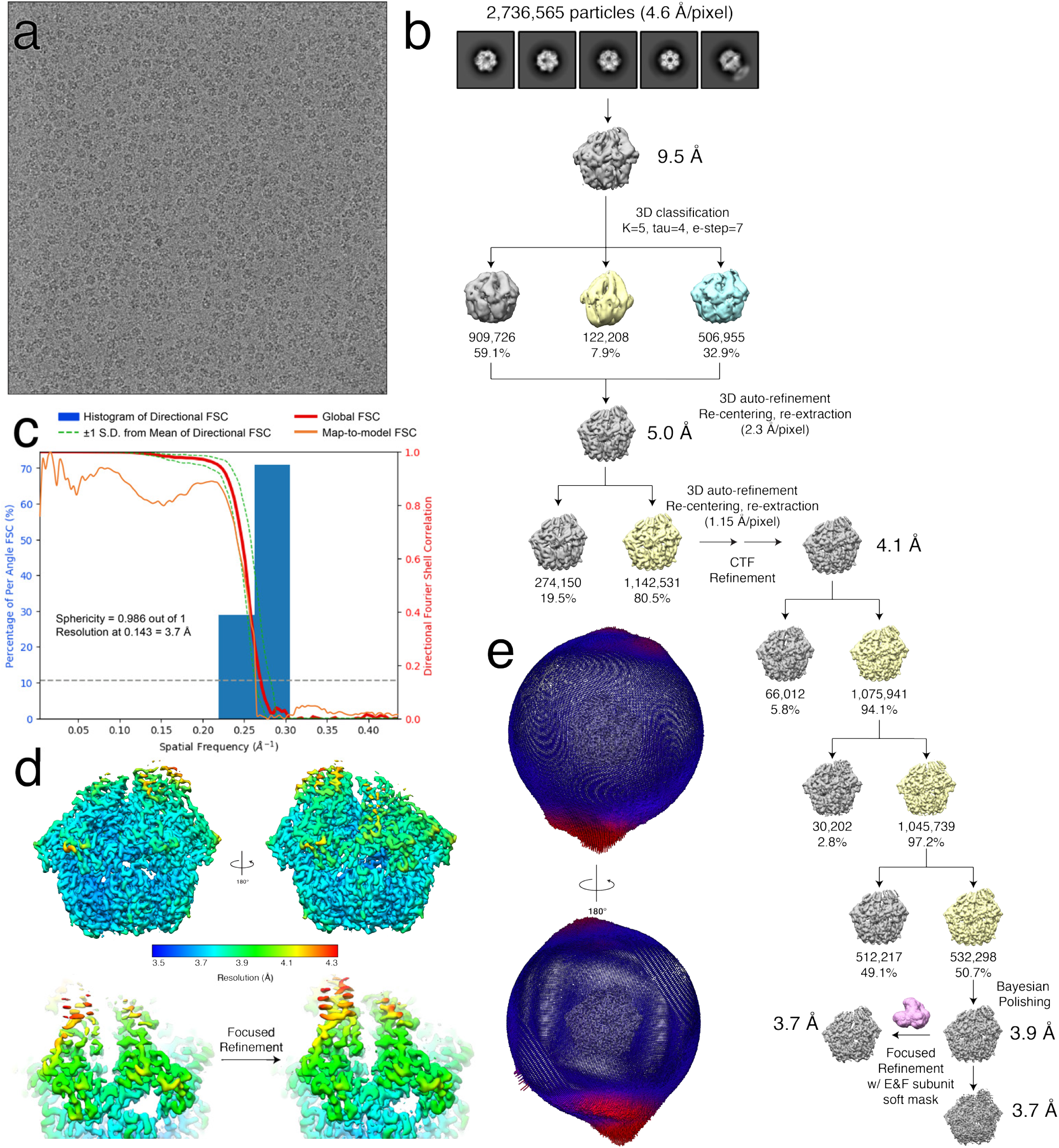
Cryo-EM structure determination of the human LONP1 bound to bortezomib. **a.** Representative micrograph from cryo-EM data collection. **b.** Cryo-EM data processing scheme followed using RELION 3.1 software^54^ to obtain the final 3D reconstruction of substrate-bound human LONP1^Bz^. Final steps included performing focused refinement with a soft mask over the ADP1 and ADP2 subunits to improve resolution of the seam subunits. The focused refinement map was used for atomic model building and refinement. **c.** 3DFSC^62^ of the final reconstruction of LONP1^Bz^ reporting a global resolution of 3.7 Å at FSC=0.143 and Sphericity of 0.986 out of 1. **d.** Final LONP1^Bz^ reconstruction colored by local resolution calculated using RELION, showing regions resolved to 3.5 Å at the core of the complex to worse than 4.1 Å in flexible regions. Focused refinement with a mask around ADP1 and ADP2 qualitatively improved the structural detail of these subunits, shown below. **e.** Euler angle distribution plot of the 532,298 particles used in the final reconstruction of the LONP1^Bz^ structure.

**Supplementary Figure 10.**
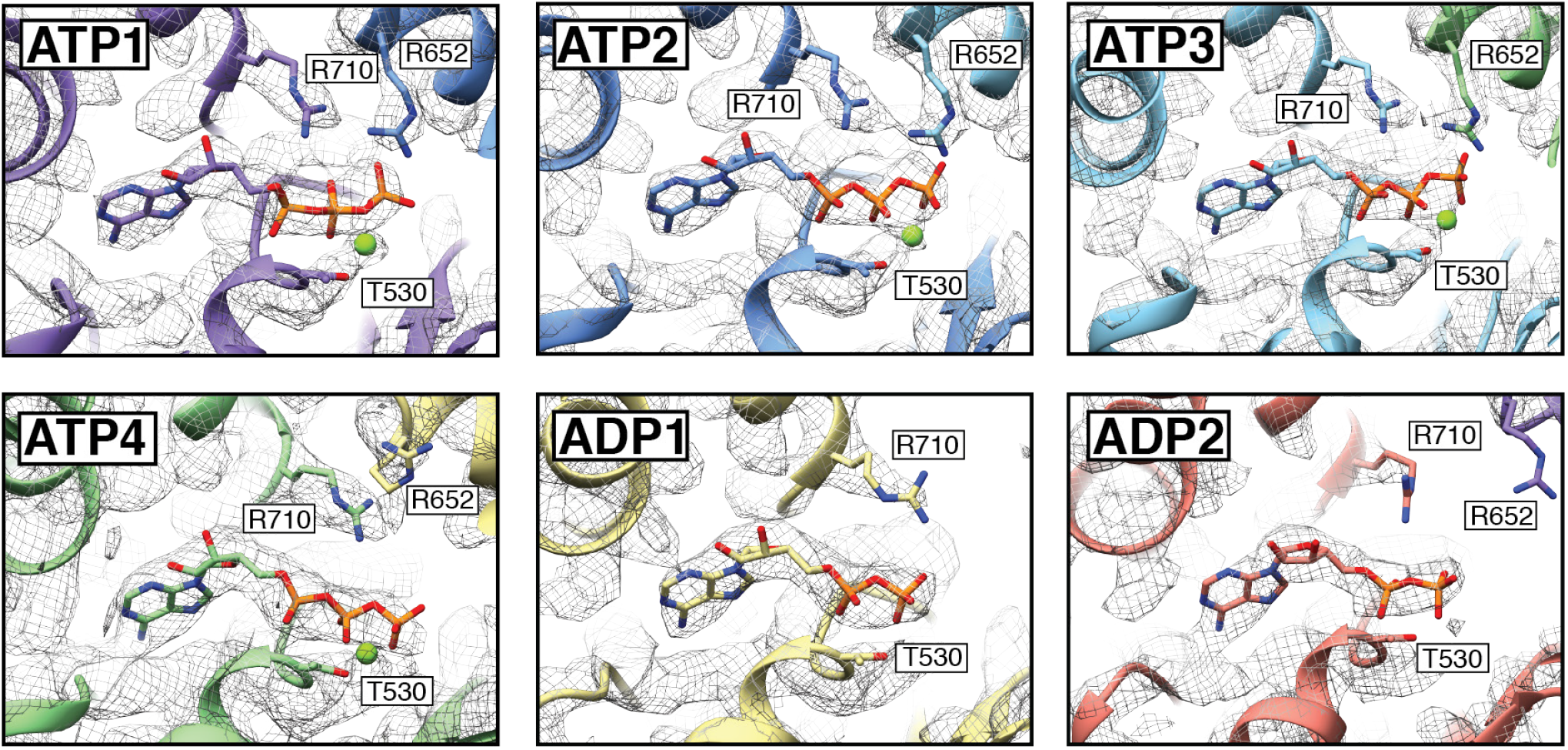
LONP1 bound to substrate and bortezomib shows distinct nucleotide densities in the nucleotide-binding pocket. The cryo-EM density, shown as an isosurface mesh contoured at sigma=0.01, in the vicinity of the nucleotide binding pocket was of sufficient quality to unambiguously assign the nucleotide state of each of the subunits. ATP1, ATP2, ATP3, and ATP4 subunits correspond to the ATPγS used to determine this structure coordinated by a magnesium cofactor. The nucleotide density in the ADP1 and ADP2 subunits corresponded to ADP molecules, as there is no apparent gamma density or magnesium.

**Supplementary Figure 11.**
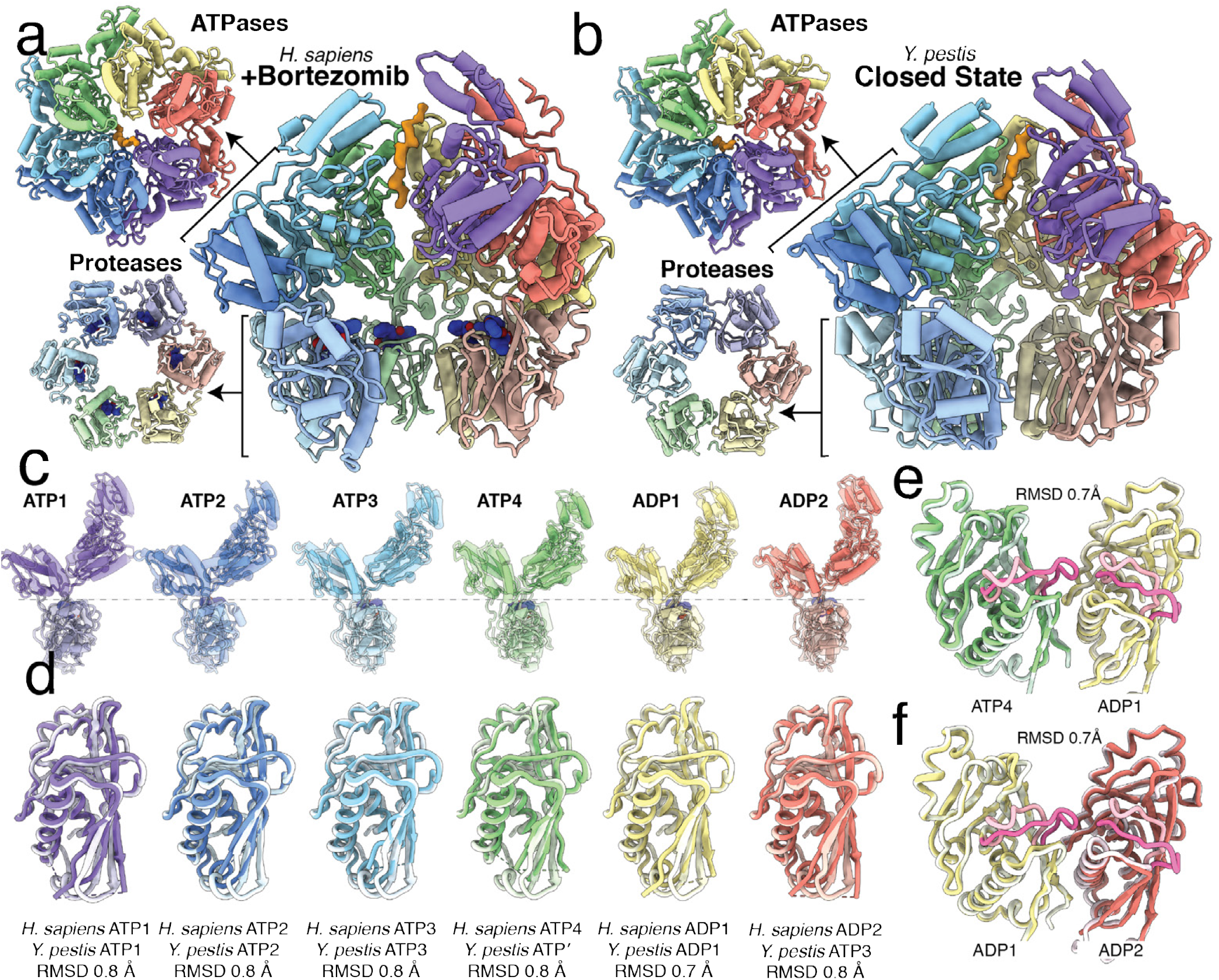
Substrate-bound LONP1^Bz^ and substrate-bound *Y. pestis* Lon protease both adopt similar proteolytically active conformations. **a,b.** Axial and lateral views of the LONP1^Bz^ atomic model (**a**) and the *Y. pestis* substrate-bound state atomic model^28^ (**b**). **c.** Individual protomers of LONP1^Bz^ aligned side-by-side relative to the protease domain produced by orienting all the proteases to a common view. The respective subunits from substrate-bound bacterial Lon are shown using a transparent ribbon representation overlaid on the subunits of substrate-bound LONP1^Bz^. In both cases, the spiraling ATPases sit atop a planar proteolytic ring. **d.** Protease domains from the LONP1^Bz^ conformer are aligned to their corresponding subunits from substrate-bound *Y. pestis* Lon. All six subunits of the protease domain are in similar conformations, with RMSDs between 0.7-0.8 Å. The protease domains of bortezomib-bound LONP1^Bz^ are activated by extending the catalytic loop, allowing catalytic dyad formation and formation of a substrate-binding groove similar to substrate-bound *Y. pestis* Lon. **e,f**. Secondary structure-based alignments between protease subunit pairs, ATP4 and ADP1 (**e**), and ADP1 and ADP2 (**f**), from substrate-bound LONP1 and LONP1^Bz^ are shown using a ribbon representation. Although the RMSDs for the subunit pair alignment are both 0.7 A, the catalytic serine-containing loop (highlighted in pink) is divergent between the substrate-bound LONP1 and LONP1^Bz^ due to the proteases adopting auto-inhibited (light pink) and activated (dark pink) conformations, respectively.

**Supplementary Figure 12.**
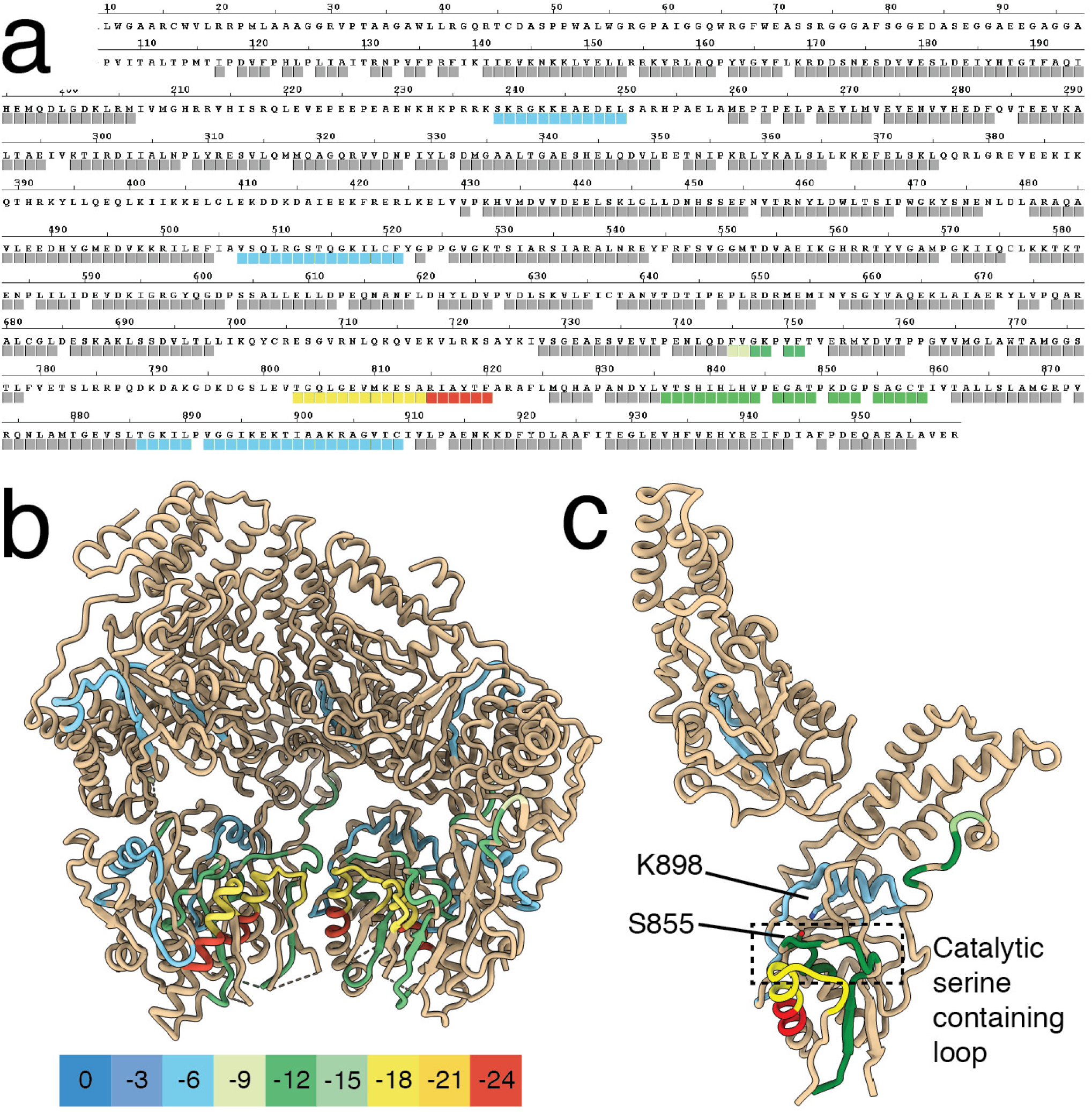
Hydrogen-deuterium exchange mass spectrometry shows a conformational change in the protease domains upon binding bortezomib. **a.** Amino acid sequence of human mitochondrial Lon. Colored boxes located underneath the amino acid sequence denote digested peptides detected in mass spectrometry experiments. Regions with no change in exchange upon the addition of Bortezomib have grey boxes while regions with reduced D_2_O exchange upon the addition of Bortezomib are colored from blue to red based on the scale shown in panel **b**; D_2_O exchange decreased in regions colored blue while regions colored in red depict where the greatest reduction in D_2_O exchange occurred. Regions with most D_2_O exchange included residues 803-814 and 815-820 colored yellow and red, respectively, belong to an alpha helix and flexible loop framing the catalytic loop the activated conformation of the protease active site. These regions likely experience a reduction in D_2_O exchange due to protease symmetrization and ring closure upon the addition of Bortezomib. **b.** Peptides that were detected as having a change in D_2_O exchange rate during hydrogen-deuterium exchange mass spectrometry are colored using the shown scale. **c.** An isolated subunit from (**b**) with a hatched box highlighting the catalytic serine-containing loop and labels for catalytic residues S855 and K898 showing reduction in D_2_O exchange in proximity of the proteolytic active site in the presence of Bortezomib.

**Supplementary Figure 13.**
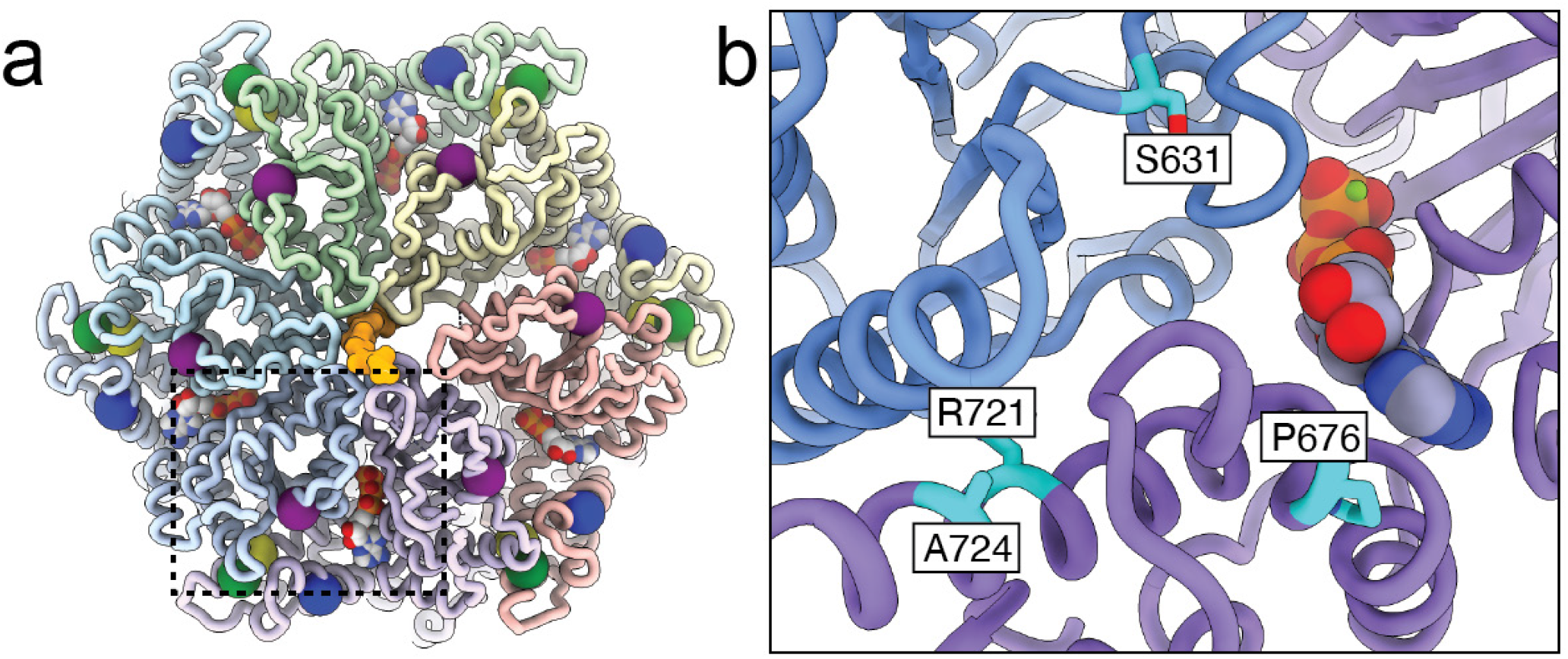
Mutations associated with CODAS syndrome localize to inter-subunit interfaces. **a.** The locations of the following point mutations associated with CODAS syndrome are denoted on the structure of the substrate-bound human LONP1 with spheres: p.Ser631Tyr (purple), p.Pro676Ser (dark blue), p.Arg721Gly (light blue), and p.Ala724Val (green). **b.** A close-up of the nucleotide binding pocket formed between ATP1 (purple) and ATP2 (blue) subunits. Residues bearing mutations associated with CODAS Syndrome (S631, P676, R721, and A724) are located at the inter-subunit interfaces, potentially playing a role in nucleotide binding, hydrolysis, or associated allostery. These mutations likely perturb inter-subunit interactions required for ATP-driven substrate translocation.

**Table S1.**
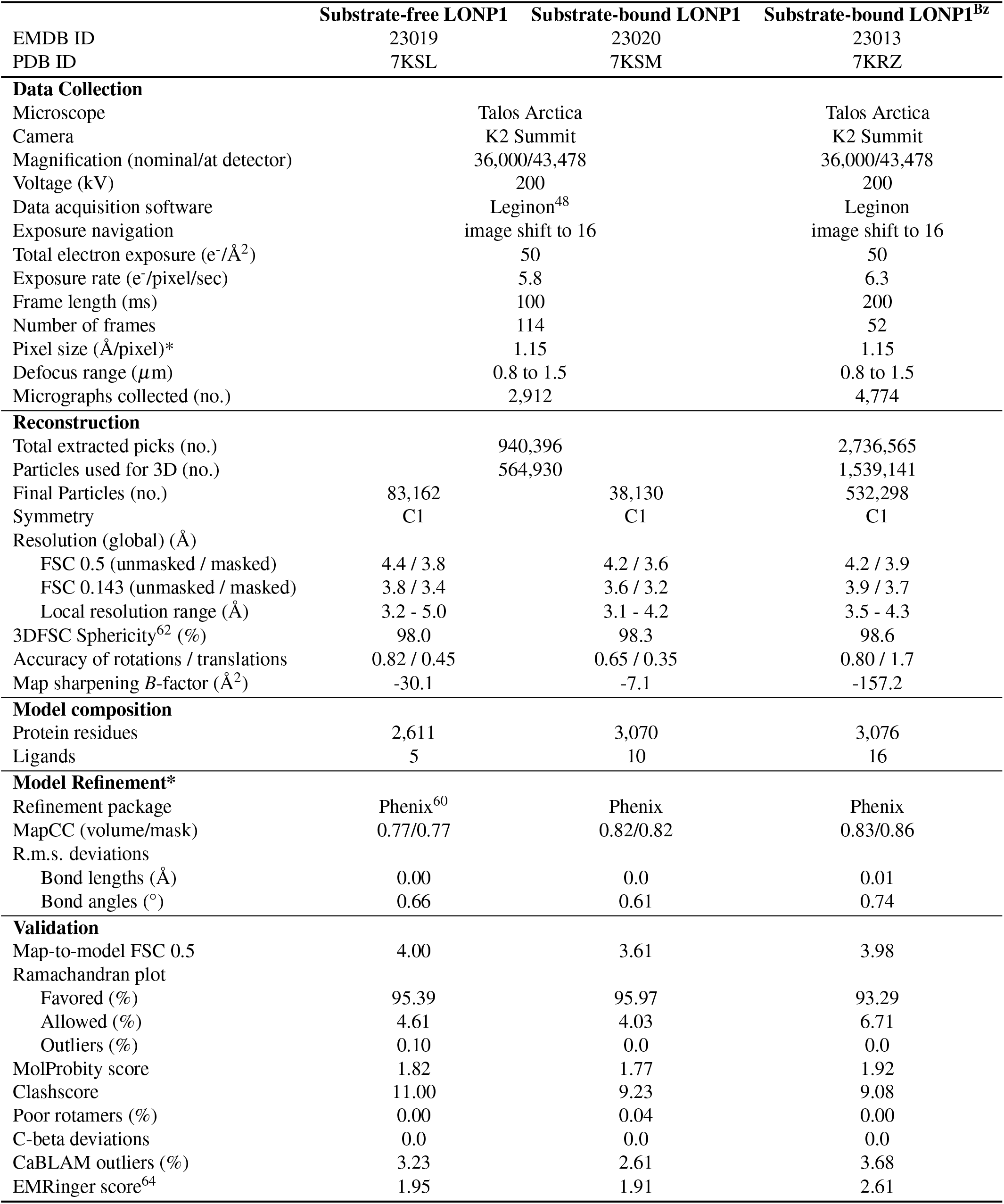
Cryo-EM data collection, refinement, and validation statistics

